# Single-cell transcriptome profiling reveals multicellular ecosystem of nucleus pulposus during degeneration progression

**DOI:** 10.1101/2021.05.25.445620

**Authors:** Ji Tu, Wentian Li, Sidong Yang, Pengyi Yang, Qi Yan, Shenyu Wang, Kaitao Lai, Xupeng Bai, Cenhao Wu, Wenyuan Ding, Justin Cooper-White, Ashish Diwan, Cao Yang, Huilin Yang, Jun Zou

## Abstract

Degeneration of the nucleus pulposus (NP) is a major contributor to intervertebral disc degeneration (IVDD) and low back pain. However, the underlying molecular complexity and cellular heterogeneity remain poorly understood. Here, we first reported a comprehensive single-cell resolution transcriptional landscape of human NP. Six novel human nucleus pulposus cell (NPCs) populations were identified by distinct molecular signatures. The potential functional differences among NPC subpopulations were analyzed at the single-cell level. Predictive genes, transcriptional factors, and signal pathways with respect to degeneration grades were analyzed. We reported that fibroNPCs, one of our identified subpopulations, might be a population for NP regeneration. CD90+NPCs were observed to be progenitor cells in degenerative NP tissues. NP-infiltrating immune cells comprise a previously unrecognized diversity of cell types, including granulocytic myeloid-derived suppressor cells (G-MDSCs). We uncovered CD11b, OLR1, and CD24 as surface markers of NP-derived G-MDSCs. The G-MDSCs were also found to be enriched in mildly degenerated (grade I and II) NP tissues compared to severely degenerated (grade III and IV) NP tissues. Their immunosuppressive function and protective effects for NPCs were revealed. Collectively, this study revealed the NPC type complexity and phenotypic characteristics in NP, providing new insights and clues for IVDD treatment.

## Introduction

Low back pain (LBP) is a major disabling health condition, with a lifetime prevalence of as high as 84% ^1^. The socio-economic burden of LBP is enormous, estimated at approximately $85 billion in 2008, and still increasing in the last decade ^2, 3^. Intervertebral disc degeneration (IVDD) is a widely recognized contributor to LBP ^4^. The degeneration of the nucleus pulposus (NP), the central gel-like part of the intervertebral disc, is a significant mechanism of IVDD ^5^. The current treatments are limited to relieving back or leg symptoms and do not focus on replenishing the NP loss and restoring the native disk structure, leading to unsatisfactory outcomes such as recrudescence or degeneration of adjacent motion segments ^6, 7^. Therefore, new therapeutic targets are urgently needed.

Nucleus pulposus cells (NPCs) are the main cell type residing in the NP, and they are responsible for maintaining tissue homeostasis ^8^. NPCs proliferate slowly and lack self-regeneration capacity adding to the intractability of the disease. Current studies on the pathophysiology of NPCs are usually supported by transcriptomic and epigenomic analyses. However, bulk-tissue level resolution masks the complexity of alterations across cells and within cell types. The uncharacterized cell types and markers residing in NP raise interest in terms of unexplored cellular heterogeneity.

The intervertebral disc (IVD) is the largest avascular organ of the body. Studies addressing immune reactions against the NP have been limited and focused on their detrimental aspects. However, given the complexity of immunity in other immune-privileged sites ^9^, immune cell subpopulations may also help restore IVD structure and lessen degeneration. Moreover, intrinsic properties of NPCs, including expression of immunomodulatory factors, and extrinsic microenvironmental changes to immune compartments, remain largely unknown. These issues highlight the urgency to understand the immune panorama of the NP during IVDD pathogenesis.

Single-cell RNA sequencing (scRNA-seq) provides a powerful alternative to study the cellular heterogeneity of NP tissues. Here, we aimed to provide the first single-cell view of IVDD pathology, profiling 39,732 cells from NP tissues across eight individuals with different grades of progressive degeneration. Notably, we comprehensively characterized the transcriptome feature of NPCs and immune cells, and we decoded the cell percentage, the heterogeneity of cell subtypes during degeneration, providing a unique cellular-level insight into transcriptional alterations associated with IVDD pathology.

## Materials and Methods

### NP tissues specimens

This study protocol was approved by the Ethics Committee of the first affiliated hospital of Soochow University. All participants signed a written informed consent form. The enrolled subjects were patients who required discectomy and/or interbody fusion for burst fracture (n=2) lumbar disc herniation or spondylolysis (n=6). The patients’ characteristics were listed in Table S1. In this study, the grading of IVDD is slightly different from Pfirrmann grading system. It is difficult to obtain Pfirrmann grade I disc tissues. Thus, our grade I-IV is Pfirrmann grade II-V accordingly. (Fig. S1A)

A standard surgical protocol assured correct sample acquisition and thus accurate selection of AF and NP tissue by experienced surgeons. For burst fracture cases, we harvested the NP tissue from the disc attached to the intact bottom endplate. For severe degenerative discs, we harvested the tissue from the central region of the NP to assure no AF tissues were harvested. If local bleeding happened during the tissue harvested, the tissues were discarded to ensure no contaminated blood in collected tissues.

We processed tissues immediately following surgical procurement, and generated single-cell suspensions within ∼45 minutes with an experimental protocol optimized to reduce artifactual transcriptional changes introduced by disaggregation, temperature, or time. Human NP tissue samples were transported in MACS Tissue Storage Solution (Miltenyi Biotec) immediately after surgical resection. After tissues were harvested, the specimens were washed with PBS for three times and further checked under a dissecting microscope to guarantee there was no contamination of blood, AF tissues or other tissues. For few cases which was difficult to distinguish between the NP and inner AF, histological staining were used to confirm the absence of AF tissues or other tissues. (Fig. S1B-D)

### Single-cell Dissociation of human nucleus pulposus

The tissues were surgically removed and kept in MACS Tissue Storage Solution (Miltenyi Biotec) until processing. The tissue samples were processed as described below. Briefly, samples were first washed with phosphate-buffered saline (PBS), minced into small pieces (approximately 1mm^3^) on ice and enzymatically digested with 500 U/mL collagenase I, 150 U/mL collagenase II, 50 U/mL collagenase IV, 0.1 mg/ml hyaluronidase, 30 U/mL DNaseI and 5% Fetal Bovine Serum Oringin South America (Yeasen) for 95 min at 37°C, with agitation. After digestion, samples were sieved through a 70 µm cell strainer, and centrifuged at 300 g for 5 min. After washing with PBS containing 0.04% BSA, the cell pellets were re-suspended in PBS containing 0.04% BSA and re-filtered through a 35 μm cell strainer. Dissociated single cells were then stained with AO/PI for viability assessment using Countstar Fluorescence Cell Analyzer. The single-cell suspension was further enriched with a MACS dead cell removal kit (Miltenyi Biotec).

### Single-cell RNA Sequencing

BD Rhapsody system was used to capture the transcriptomic information of the (8 sample-derived) single cells. Single-cell capture was achieved by random distribution of a single-cell suspension across >200,000 microwells through a limited dilution approach. Beads with oligonucleotide barcodes were added to saturation so that a bead was paired with a cell in a microwell. Cell-lysis buffer was added so that poly-adenylated RNA molecules hybridized to the beads. Beads were collected into a single tube for reverse transcription. Upon cDNA synthesis, each cDNA molecule was tagged on the 5′ end (that is, the 3′ end of a mRNA transcript) with a unique molecular identifier (UMI) and cell label indicating its cell of origin. Whole transcriptome libraries were prepared using the BD Rhapsody single-cell whole-transcriptome amplification workflow. In brief, second strand cDNA was synthesized, followed by ligation of the WTA adaptor for universal amplification. Eighteen cycles of PCR were used to amplify the adaptor-ligated cDNA products. Sequencing libraries were prepared using random priming PCR of the whole-transcriptome amplification products to enrich the 3′ end of the transcripts linked with the cell label and UMI. Sequencing libraries were quantified using a High Sensitivity DNA chip (Agilent) on a Bioanalyzer 2200 and the Qubit High Sensitivity DNA assay (Thermo Fisher Scientific). The library for each sample was sequenced by illumina sequencer (Illumina, San Diego, CA) on a 150 bp paired-end run.

### Single-cell RNA Statistical Analysis

We applied fastp^10^ with default parameter filtering the adaptor sequence and removed the low quality reads to achieve the clean data. UMI-tools was applied for Single Cell Transcriptome Analysis to identify the cell barcode whitelist. The UMI-based clean data was mapped to human genome (Ensemble version 91) utilizing STAR mapping with customized parameter from UMI-tools standard pipeline to obtain the UMIs counts of each sample. Cells contained over 200 expressed genes and mitochondria UMI rate below 20% passed the cell quality filtering and mitochondria genes were removed in the expression table. Seurat packageSeurat package (version: 2.3.4, https://satijalab.org/seurat/) was used for cell normalization and regression based on the expression table according to the UMI counts of each sample and percent of mitochondria rate to obtain the scaled data.In order to correct the batch effect, fastMNN function from scater package (https://github.com/Alanocallaghan/scater/tree/master/R) was applied with k value equals 5 and UMAP as well as tSNE dimension reduction construction was calculated following the batch effect correction result. PCA was constructed based on the scaled data with top 2000 high variable genes and top 10 principals were used for tSNE construction and UMAP construction.

Utilizing graph-based cluster method (resolution = 0.8), we acquired the unsupervised cell cluster result based the PCA top 10 principal and we calculated the marker genes by FindAllMarkers function with wilcox rank sum test algorithm under following criteria:1. lnFC > 0.25; 2. pvalue<0.05; 3. min.pct>0.1. In order to identify the cell type detailed, the clusters of same cell type were selected for re-tSNE analysis, graph-based clustering and marker analysis. To identify differentially expressed genes among samples, the function FindMarkers with wilcox rank sum test algorithm was used under following criteria:1. lnFC > 0.25; 2. pvalue<0.05; 3. min.pct>0.1.

### Pseudo-Time Analysis

We applied the Single-Cell Trajectories analysis utilizing Monocle2 (http://cole-trapnell-lab.github.io/monocle-release) using DDR-Tree and default parameter. Before Monocle analysis, we select marker genes of the Seurat clustering result and raw expression counts of the cell passed filtering. Based on the pseudo-time analysis, branch expression analysis modeling (BEAM Analysis) was applied for branch fate determined gene analysis.

More Single-Cell Trajectories analysis was applied by utilizing PAGA in scanpy package (https://scanpy.readthedocs.io/en/latest/index.html, version 1.6.0) and Slingshot (https://bioconductor.org/packages/release/bioc/html/slingshot.html version 1.4.0). Before analysis, we select marker genes of the Seurat clustering result and raw expression counts of the cell passed filtering.

### Cell Communication Analysis and SCENIC Analysis

To enable a systematic analysis of cell–cell communication molecules, we applied cell communication analysis based on the CellPhoneDB, a public repository of ligands, receptors and their interactions. Membrane, secreted and peripheral proteins of the cluster of different time point was annotated. Significant mean and Cell Communication significance (p-value<0.05) was calculated based on the interaction and the normalized cell matrix achieved by Seurat Normalization. To assess transcription factor regulation strength, we applied the Single-cell regulatory network inference and clustering (pySCENIC, v0.9.5) workflow, using the 20-thousand motifs database for RcisTarget and GRNboost.

### QuSAGE Analysis (Gene Enrichment Analysis)

To characterize the relative activation of a given gene set such as pathway activation, “CellDeath_Ferrdb_soring” and “CellDeath_Inflammasome_review” as described before, we performed QuSAGE (2.16.1) ^11^ analysis.

### Cell fate analysis including CytoTRACE and Velocity

We applied the scVelo package with default parameter for studying the cellular differentiation status based on the bam mapping file from UMI tools STAR mapping steps to solve the transcriptional dynamics of splicing kinetics using a likelihood-based dynamical model. We applied CytoTRACE Analysis for cell stem stage analysis with default parameter. ^12^

### Pathway Analysis

Pathway analysis was used to find out the significant pathway of the marker genes and differentially expressed genes according to KEGG database. We turn to the Fisher’s exact test to select the significant pathway, and the threshold of significance was defined by P-value and FDR.

### Rat model of IVDD

A total of 15 male Sprague–Dawley rats, aged 3 months, were used for the experiments in vivo. Ten rats underwent the surgeries, and the remaining five rats underwent no surgical intervention as negative controls. On the day of surgery, the animals were anaesthetized in the induction chamber with oxygen flow at 1 liter per minute and up to 4% isoflurane. The isoflurane was reduced to between 2 and 2.5% once the animal is asleep. Once anesthetized, the animal was placed on a heating pad in supine position. Buprenorphine (0.1 mg/kg) in saline were administered by subcutaneous injection for post operation pain relief. The intervertebral space was located by digital palpation. Next a needle was affixed using a clamp so that 5mm tip sticks out. After cleaned injection site with ethanol, the needle head then be inserted into the intervertebral space and held in place for 20 sec to ensure puncture. After surgeries, the animals were returned to a warm and clean cage and monitored until wake.

### Quantitative real-time PCR

The total RNA was harvested from the NPCs and it was reverse transcribed into cDNA with SuperScript II reverse transcriptase (Invitrogen, Cat No. 18064014). The sequence of primers for RT-PCR are as below: MSMO1 (Forward 5’-3’: TGCTTTGGTTGTGCAGTCATT; Reverse 5’-3’: GGATGTGCATATTCAGCTTCCA) HMGS1(Forward Primer: GAGCCCATACTCATCAAGTACCG; Reverse Primer: CCTCGGGAGAGATGCACAC); DKK (Forward Primer: ACGAGTGCATCATCGACGAG; Reverse Primer: GCAGTCCCTCTGGTTGTCAC); FN1 (Forward Primer: AGGAAGCCGAGGTTTTAACTG; Reverse Primer: AGGACGCTCATAAGTGTCACC); CRTAC1(Forward Primer: TGTCCAGGATGTTACCGTTCC; Reverse Primer: AGCTGGGTGGGATTACTGTCA); COL1A1(Forward Primer: ATCAACCGGAGGAATTTCCGT; Reverse Primer: CACCAGGACGACCAGGTTTTC); COL3A1(Forward Primer: GGAGCTGGCTACTTCTCGC; Reverse Primer: GGGAACATCCTCCTTCAACAG); MMP2 (Forward Primer: CCCACTGCGGTTTTCTCGAAT; Reverse Primer: CAAAGGGGTATCCATCGCCAT); RPS29 (Forward Primer: CGCTCTTGTCGTGTCTGTTCA; Reverse Primer: CCTTCGCGTACTGACGGAAA); RPS21 (Forward Primer: AGCAATCGCATCATCGGTG; Reverse Primer: CCCCGCAGATAGCATAAGTTTTA); CHI3L1 (Forward Primer: AAGCAACGATCACATCGACAC; Reverse Primer: TCAGGTTGGGGTTCCTGTTCT); NF-kB (Forward Primer: AACAGAGAGGATTTCGTTTCCG; Reverse Primer: TTTGACCTGAGGGTAAGACTTCT); CXCL2 (Forward Primer: CTTGTCTCAACCCCGCATC; Reverse Primer: CAGGAACAGCCACCAATAAGC); FMOD (Forward Primer: ATTGGTGGTTCCACTACCTCC; Reverse Primer: GGTAAGGCTCGTAGGTCTCATA)

### Scoring of biological processes

Individual cells were scored for their expression of gene signatures representing certain biological functions. The functional signatures were derived from the Gene Ontology database. The immune response and steroid biosynthetic process were measured by GO:0006955 and GO:0006694 respectively. The innervation score was measured by the calculating the average expression of genes in the Gene Ontology term ‘innervation’ (GO:0060384). The angiogenesis was measured by term ‘positive regulation of angiogenesis’ (GO:0045766). The indicated scores were calculated by scaling the normalized expression of a gene across all cells. Gene weights were set to either 1 or −1 to reflect positive or negative relationships.

### Immunohistochemical (IHC) assays

NP tissues were fixed for 48 h in 4% buffered paraformaldehyde. The sections were pre-treated for 10 min with trypsin (0.05%) and then treated with 3% (vol/vol) H2O2 for 15 min. Then, the sections were blocked at room temperature for 1h with 10% goat serum. After washing with PBS, sections were incubated with anti-CRTAC1 (1:50 dilution, ab25469, Abcam), anti-CHI3L1 (1:250 dilution, ab255297, Abcam), anti-MSMO1 (1:100 dilution, ab203587, Abcam), anti-RPS14 (1:50 dilution, Abcam), anti-FRZB (1:100 dilution, ab205284, Abcam), anti-MMP2 (1:200 dilution, ab86607, Abcam), antibodies overnight at 4°C.The sections were then washed with PBS and incubated with a biotinylated secondary antibody for 15 min from a Histostain Plus kit (Invitrogen, CA, USA). The sections were then washed and incubated with 3, 3’-diaminobenzidine for 2 min. Lastly, using light microscopy to observe the section.

### Cell Sorting

Tissue samples were harvested from patients with IVDD and mechanically dissociated to generate single cell suspensions as described above. Cells were blocked with FcR Blocking Reagent, human (Miltenyi, 130-059-901) on ice for at least 10 min. Cells were then centrifuged at 300g for 5 min at 4°C and washed once with BSA running buffer (0.5%BSA). Cells were incubated for 30 min at 4°C with pre-conjugated fluorescent labeled antibodies with the following combinations: CD45 (BioLegend, 368532 (PE/Cyanine7)), CD11b (BioLegend, 301330(FITC)), OLR1 (BioLegend, 358604 (PE)), CD24 (BioLegend, 311118(APC)). Cells sorted by Beckman Moflo XDP and desired populations were isolated for different experiments. For CD90+ NPCs, dissociated cells were incubated with CD90 microbeads (Miltenyi Biotec, 130-096-253; 1 μl per 107 cells) to enrich CD90+ NPCs.

### Multilineage differentiation assays in vitro

For osteogenic differentiation, the CD90+ NP cells were cultured in a humidified incubator (37°C, 5% CO_2_) with renewal of the culture medium every 3 days. The cells were incubated in a differentiation medium for 2–4 weeks once the cells had reached 100% confluence, during which time the medium was changed every 2–3 days. The differentiation medium was as follows: DMEM-LG supplemented with 10% FBS, 1 μM dexamethasone (Sigma), 50 μg/ml ascorbic acid (Sigma), 10 mM sodium β-glycerophosphate (Sigma) and 1% penicillin/streptomycin (Sigma). The cells were fixed with ice-cold 70% ethanol and stained with Alizarin Red S (Amresco, Solon, OH, USA), as well as the von Kossa stain, to detect mineralization (calcium deposits). The alkaline phosphatase (ALP) activity was also tested. For adipogenic differentiation, the CD90+ cells were first grown to 100% confluence and then incubated for 3 days in an induction medium consisting of DMEM-LG supplemented with 10% FBS, 100 μM indomethacin (Sigma), 0.1 μM dexamethasone, 0.5 mM IBMX, 10 μg/ml human insulin and 1% penicillin/streptomycin. The cells were incubated in the induction and maintenance media for >2 weeks and then fixed with 4% paraformaldehyde for 30 min at room temperature and stained with Oil Red O, as well as Sudan Black B, to detect fat deposition. For chondrognic differentiation, cells were maintained as pellet cultures (2.5–5 × 105 cells/pellet) in DMEM high-glucose medium supplemented with insulin transferin selenium (with albumin), sodium pyruvate (100 μg/mL), l-proline (40 μg/mL), ascorbate 2-phosphate (50 μg/mL), and TGF-β3 (10 ng/mL).

### Western Blotting

Western blotting was performed to analyze protein levels. The cells were lysed using RIPA Lysis Buffer, and protein concentrations were determined using the BCA Assay. Nuclear and cytoplasmic proteins were lysed using the Nuclear/Cytosolic Fractionation assay kit from BioVision (Mountain View, CA, USA) following the manufacturer’s protocol. The membrane was blocked with 5% non-fat milk and then incubated with the following primary antibodies at 4 °C overnight: anti-BAX (1:1000), anti-Bcl-2 (1:1000), anti-NLRP3 (1:1000)anti-aggrecan (1:1000), anti-MMP-13 (1:4000)After this incubation, the membranes were incubated for 2 h on a shaker at 37 °C with horseradish peroxidase (HRP)-conjugated secondary antibodies (Boster, Wuhan, China) used at a dilution of 1: 2000 and then washed. Finally, the protein bands were visualized and detected using the enhanced chemiluminescence (ECL) system, and immunoreactive bands were quantified using the ImageQuant LAS 400 software (GE Healthcare Life Sciences) and calculated by normalization to the reference bands of GAPDH or lamin B.

### Histopathology and Immunofluorescence

For histopathological analysis, tissues were fixed in 10% formalin for 96 h, decalcified in 10% ethylenediaminetetraacetic acid (EDTA) for at least one month, embedded in paraffin, and 4–5 μm sections were stained with Safranin O/ Fast Green Stain. For Immunofluorescence, the caudal disc sections were blocked in 5% normal serum (Thermo Fisher Scientific, 10000 C) in PBS-T (0.4% Triton X-100 in phosphate-buffered saline (PBS)), and incubated with the primary antibody. The primary antibodies used were CD11b (1:500, 120772, Absin), OLR1(1:500, Absin, 123947), CD24 (1:200, Proteintech, 10600-1-AP). After washed with PBS/1% BSA, the sections were incubated with Alexa-conjugated antibodies and washed with PBS/1% BSA followed by PBS. The DAPI was used for counterstaining.

### T cell Suppression Assay

T lymphocytes were isolated from healthy donor’s PBMCs via Pan T Cell Isolation Kit (Miltenyi Biotec,130-096-535) following the manufacturer’s instructions. Isolated T cells were then cultured in 96-well round-bottom plates in complete culture medium containing soluble, or plate-bound, anti-CD3 (1 μg/ml) and soluble anti-CD28 (5 μg/ml). Sorted CD45+CD11b+OLR1+CD24+ and CD45+CD11b+OLR1+CD24-cells were added to T cells in 1:1 ratio. After 4 days of culture, T cell proliferation was assessed by MTT assay. T cells without sorted cells co-cultured was used as positive control.

### ROS Production Assay

After isolating NP-derived MDSCs by fluorescence-Activated Cell Sorting, the cellular ROS level was determined with the ROS assay kit. The cell suspension was used at a concentration of 1 × 10^6^ cells/ml and was diluted with DCFH-DA solution (10 µM) and incubated at 37 °C for 20 min with mixing by inversion every 5 min. After washing the cells with serum-free medium, the sample was analyzed by flow cytometry (FAC Scan, Becton Dickenson, USA). Rosup was used as positive control.

## Results

### Single-cell profiling of NP in human subjects with IVDD pathology

We dissociated NP tissues from eight IVDD patients (TableS1) with different degeneration degrees (see method) and performed scRNA-seq on the BD Rhapsody system (Fig. 1A, Fig. S1A). After quality control filtering to remove cells with low gene detection (<600 genes) and high mitochondrial gene content (>8%) (Fig. S1E-H), we identified and annotated the major cell types by evaluating the expression of specific known genes (Fig.1B-C). NPCs were identified by SOX9 and ACAN. Natural killer (NK) cells were identified by CD94 ^13^. Macrophage/monocytes were identified by CD163^14^, and T cells were identified by TRAC (T Cell Receptor Alpha Constant). (Fig. 1B-C).

**Figure 1.**
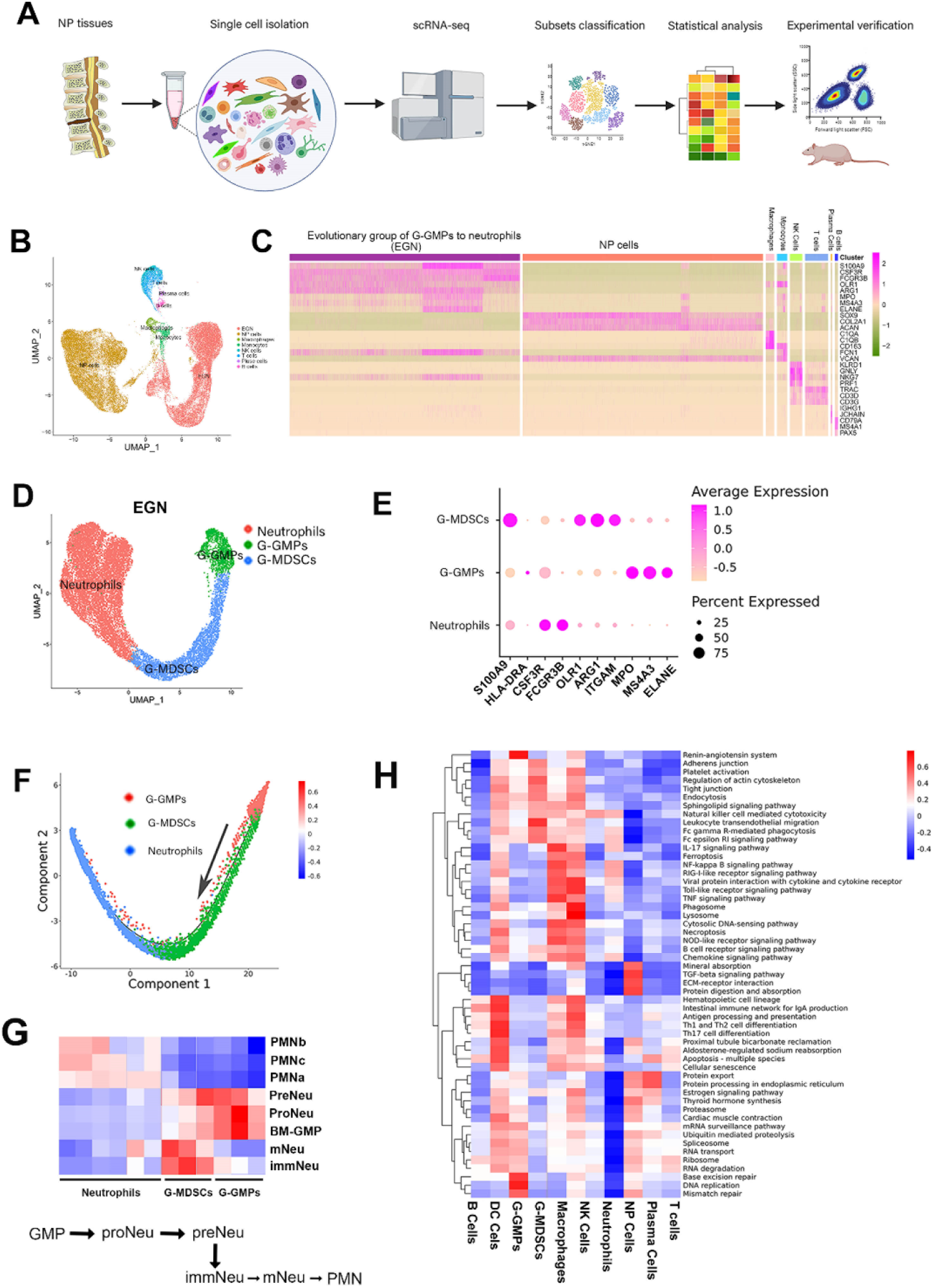
Single-cell profiling of nucleus pulposus (NP) in human subjects with intervertebral disc degeneration (IVDD) pathology. **(A)** Graphical representation of the experimental workflow. **(B)** Uniform Manifold Approximation and Projection (UMAP) visualization showing all annotated cells. **(C)** Heatmap showing the typically expressed genes in each cell type. **(D)** UMAP visualization showing GMP, granulocytic myeloid-derived suppressor cell (G-MDSCs), and neutrophils. **(E)** Dot plot showing scaled expression of selected signature genes for GMP, G-MDSCs, and neutrophils, by average expression of each gene in each cluster scaled across all clusters. Dot size represents the percentage of cells in each cluster with more than one read of the corresponding gene. **(F)** Monocle method reconstructed pseudospace trajectory for GMP, G-MDSC, and Neutrophils. **(G)** Correlation of our defined neutrophils, G-MDSCs, and G-GMPs with the neutrophil subtypes reported by previous single-cell publications ^22^. **(H)** QuSAGE analysis was used to identify pathways with significantly (FDR<0.005) altered activity. Coloring represents down-regulated (blue) to up-regulated (red)activity relative to the other cell types. (FDR < 0.1).

Apart from the cell types described above, we also noted a group of cells transcriptionally clustered together and are hard to annotate, as they expressed the markers of three immune cell types, including granulocyte–monocyte progenitors (G-GMPs), granulocytic myeloid-derived suppressor cell (G-MDSC), as well as neutrophils. To distinguish these three cell types, we further clustered this cell group and produced 7 transcriptionally distinct pre-clusters (Fig. S2A-C). Based on known gene markers, we identified G-GMPs (expressing MS4A3, MPO, and ELANE) ^15, 16^, neutrophils (FCGR3B (CD16b) + HLADR) ^17^, and granulocytic myeloid-derived suppressor cell (G-MDSC, ITGAM (CD11b), OLR1, and ARG1) ^18, 19^. (Fig.1D-E) Monocle trajectory analysis revealed GMPs were distributed at the start of the trajectory, G-MDSCs at the intermediate segment, and neutrophils were dispersed along the trajectory (Fig. 1F). Thus, we referred to them as an evolutionary group of G-GMPs to neutrophils (EGN).

Next, we compared our identified G-GMPs, G-MDSCs, and neutrophils with those reported in previous publications and single-cell data (Fig. 1G) ^20, 21^. G-GMPs were correlated with Bone marrow (BM)-GMP, pre-neutrophils (preNeu), and neutrophil progenitors (proNeu). G-MDSCs were correlated with immature neutrophils (immNeu) and mature neutrophils (mNeu).Neutrophils were correlated with polymorphonuclear morphology neutrophils (PMN). It has been previously reported that dynamic transitions between neutrophil subpopulations exist: PMN cells can arise from both mNeu and immNeu, and GMP can be differentiated into proNeu and preNeu (Fig.1G) ^22^. Taken together, our data suggest that G-GMPs are responsible for the generation of neutrophils, and G-MDSCs. The latter is a putative intermediate differentiated status.

We next explored signal pathways using gene set variation analysis of all cell types in NP tissues. TGF-beta, extracellular matrix (ECM)-receptor interaction, and protein metabolism were upregulated in NPCs; chemokines, FcR1, and NF-κB were upregulated in neutrophils. In G-GMPs, base excision repair and DNA replication were upregulated. Phagocytosis and transendothelial migration were upregulated in G-MDSCs. The Toll-like pathway and antigen presentation were upregulated in macrophages (Fig. 1H).

### Identification of human NPC atlas

Consistent with previous studies ^23, 24^, we found a lack of distinctive clusters in the tSNE map (Fig. 2A) which may suggest the heterogeneity exhibited from these cell populations. Six subpopulations were identified based on their highly expressed genes and published single-cell studies (Fig. 2B):

**Figure 2.**
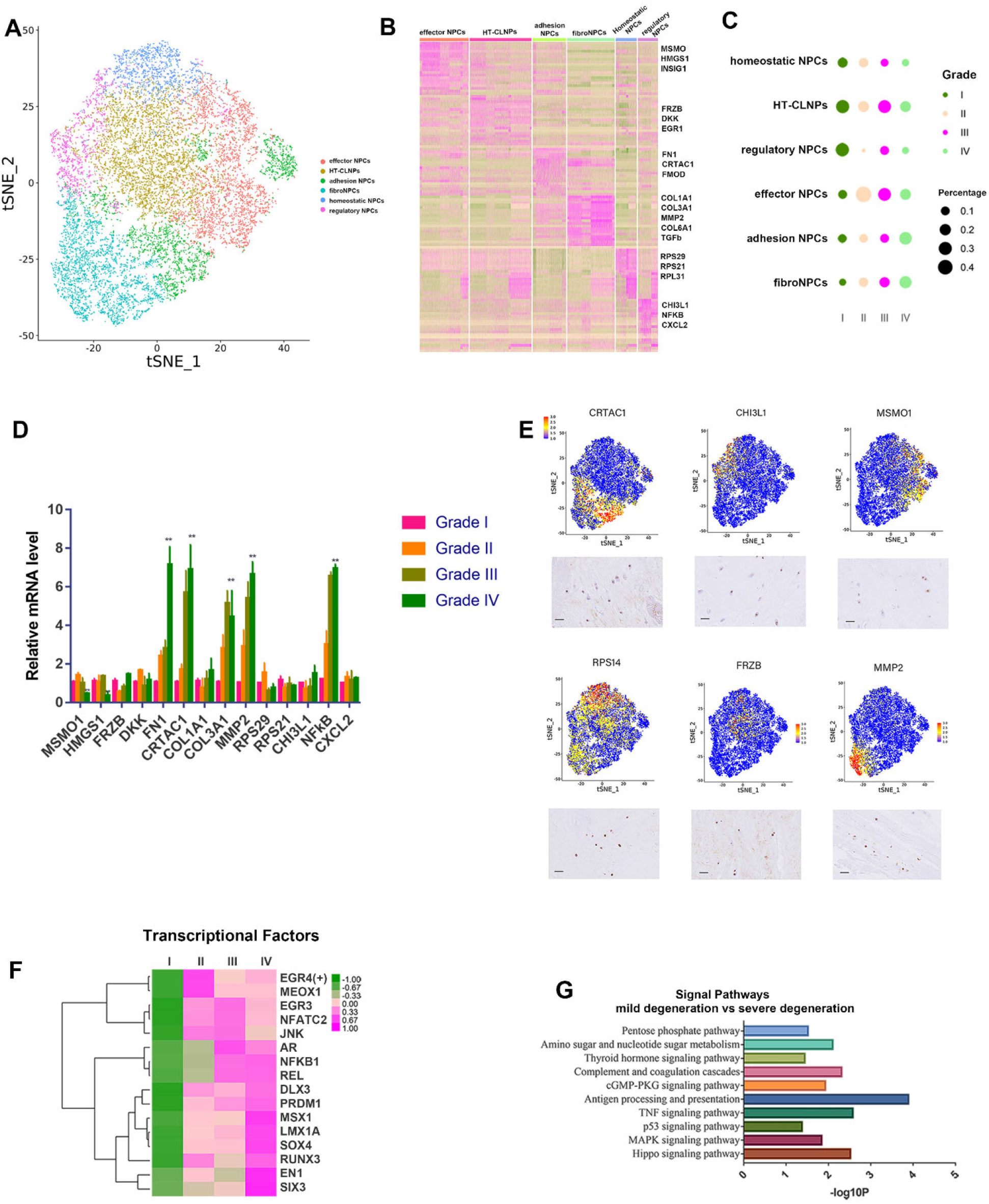
Identification of human NPC atlas and transcriptional changes correlated with IVDD severity. **(A)** UMAP visualization of human NP cells identified six different clusters after unsupervised clustering. **(B)** Heatmap revealing the scaled expression of differentially expressed genes for each cluster. **(C)** Dot plots showing the grade distribution in each NP cell subsets. **(D)** RT-qPCR for the representative genes of NPC atlas in different degenerative grades discs. (n = 3 with mean ± SD shown) **(E)** Representative immunohistochemistry assay of indicated genes in NP tissues. **(F)** Heatmap showing grade-related transcription factors. **(G)** Enriched signal pathways related with degeneration grades.

1. hypertrophy chondrocyte-like NPCs (HT-CLNPs; Groups 0, 1; expressing FRZB, DKK) ^25, 26^;
2. effector NPCs (Groups 2, 4; expressing genes related with metabolism, like MSMO1^27^, HMGCS1^28^);
3. homeostatic NPCs (Group 5; expressing RPS29 and RPS21) ^24^;
4. regulatory NPCs (Group 6; expressing CHI3L1 ^24^, CXCL2, and NFKB ^29^);
5. fibroNPCs (Groups 8, 9, and 10, expressing genes related with fibrosis, COL1A1, COL3A1, COL6A1) ^23, 24^;
6. adhesion NPCs (Groups 3, 11; expressing genes related to cell adhesion and migration, such as FN1 ^30^ and CRTAC1^31^).

We next analyzed the relationship between the degeneration grades and distributions of the cell populations. HT-CLNPs and regulatory NPCs were the main cells for grade I discs. Effector NPCs and HT-CLNPs were mainly subpopulations in grades II and III discs. For grade IV discs, fibroNPCs and adhesion NPs were the main NPCs (Fig. 2C). Moreover, the proportion of adhesion NPCs and fibroNPCs increased with the severity of IVDD (from grade I to IV), while the proportion of homeostatic NPCs showed opposite trends.

Real-time quantitative PCR (qPCR) was used to validate the indicated gene expression in NPCs from different degeneration grades. The results revealed that the expressions of FN1 and CRTAC1, COL3A1, and MMP2 (markers of adhesion NPCs and fibroNPCs) were significantly elevated in NPCs from late-stage degenerative discs (grade III and IV). The expressions of homeostatic NPC markers, MSMO1 and HMGS1, significantly decreased in NPCs from late-stage degenerative discs. (Fig. 2D). We then performed an immunohistochemistry assay to validate the expressions of markers for each NPC subpopulation at the protein level. (Fig. 2E).

### Identification of NPC transcriptional changes correlated with IVDD severity

A group of transcriptional factors (TFs) were idnetifed to be related to grade of degeneration.(Fig.2F) Some TFs, like AR, REL, LMX1A, and PRDM1, were first revealed. (Fig. 2F). Enrichment for TFs of REL in grade III and IV discs may be related to Rel/NF-κB signal transduction pathway in IVDD ^32^.

Besides, we compared differentially expressed genes (DEGs) between severe (grades III and IV) and mild (grades I and II) degeneration and identified candidate genes related to IVDD progression, including HSPH1, CTGF, MMP13, and HSP90AA1 (Table S2). There are significantly elevated expressions of a list of heat shock protein (HSP) genes in severe degenerative discs, including HSPA1B, HSPH1, HSP90AA1, HSPA8, HSPA1A, HSPB8, and HSPD1, indicating the importance of stress-related mechanisms in IVDD. Enrichment for TF, SOX4, and INHBA genes is consistent with the importance of TGF-β signaling in IVDD ^33^. ANXA5 was also elevated, which may be related to mitochondrial dysfunction induced cell apoptosis ^34^.

Based on a IVDD grade-related gene set, a group of signaling pathways were found to potentially promote degeneration, including antigen processing and presentation, the TNF pathway, MAPK, and Hippo pathways (Fig. 2G).

### ScRNA-seq reveals transcriptional features of NPCs subpopulations

To analyze the functional differences among subpopulations, we used the Quantitative Set Analysis for Gene Expression (QuSAGE) (Fig. 3A-C) and Gene Ontology (GO) database (Table 3) ^11, 35^ to investigate the biological process of our identified NPCs subpopulations. Effector NPCs were enriched with metabolic process-related genes (e.g., sterol biosynthesis and glycosaminoglycan metabolism) and positive regulation of extracellular matrix (ECM) assembly. Regulatory NPCs were enriched with highly expressed genes responsible for cellular responses to inflammation and endogenous stimuli, such as CXCL3, IL-6, and CHI3L1, indicating that these cells might potentially regulate immune functions. Homeostatic NPCs were enriched for processes related to cellular homeostasis, including translational regulation and protein/RNA metabolism. Adhesion NPCs were enriched for cell migration and cell-matrix adhesion.

**Figure 3.**
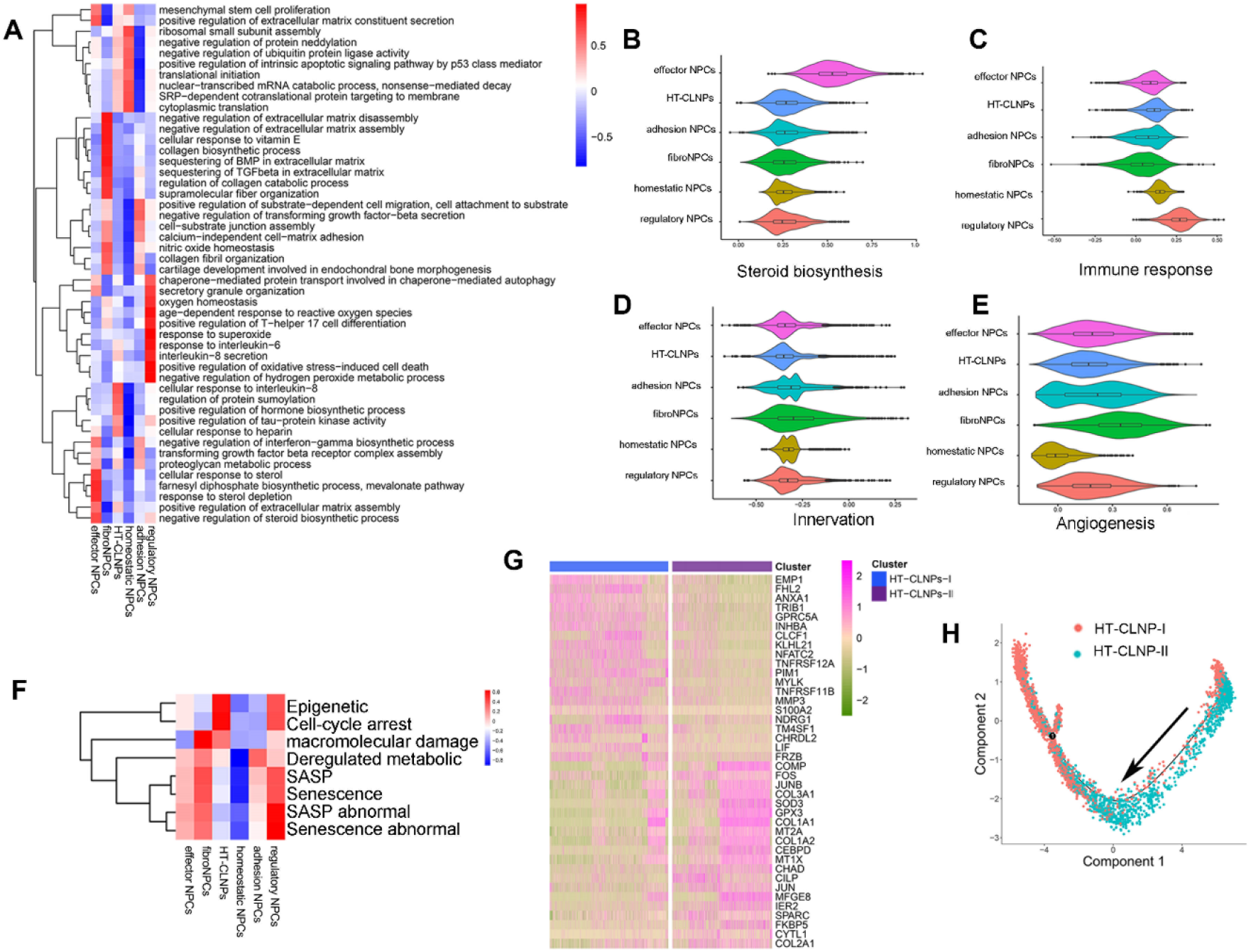
ScRNA-seq reveals transcriptional features of NPCs subpopulations. **(A)** QuSAGE analysis of cell subpopulation specific differential expression colored by statistically significant normalized enrichment scores. **(B-E)** Violin plots of steroid biosynthesis, immune response, innervation, and angiogenesis score for each cluster. **(F)** Correlation of scRNA-seq defined NPCs subpopulations with cell senescence. **(G)** Heatmap showing the scaled expression of the differentially expressed genes (DEGs) for HT-CLNP-I and HT-CLNP-II subsets. **(H)** Pseudotime trajectory axis revealing the progression of HT-CLNP-I and HT-CLNP-II.

We specifically scored for innervation, angiogenesis, and cell senescence, which are important events related to IVDD ^36, 37^, based on GO terms and published literature (see methods). It was suggested that there is no significant difference for innervation among NPC subpopulations (Fig. 3D). For angiogenesis, fibroNPCs showed the highest score, and homeostatic NPCs showed the lowest score. (Fig. 3E)

Also, it was indicated that senescence-associated secretory phenotype (SASP) was almost not expressed in HT-CLNPs or homeostatic NPCs. However, HT-CLNPs were active for cell cycle arrest (DNA damage) and SASP-related epigenetics changes. FibroNPCs were highly active for macromolecular damage, indicating that HT-CLNPs showed different cellular senescence mechanisms to FNPs (Fig. 3F).

Overall, our findings suggest differences in functions and biological processes among NPC subpopulations. Moreover, multiple potential mechanisms for cell senescence exist in different subpopulations.

Two transcriptionally distinct clusters were identified: HT-CLNP-I and HT-CLNP-II.TheDEGs of these two clusters are shown in Fig. 3G. The HT-CLNP-I was enriched in genes related to programmed cell death and negative regulation of cell proliferation. In contrast, the HT-CLNP-II subset was enriched in genes related to ECM organization and ECM disassembly. (Table S4) Additionally, the Monocle pseudotime trajectory revealed progression of the HT-CLNP-I and HT-CLNP-II clusters (Fig. 3H). These findings contribute to a deeper understanding of the hydrophilic mechanisms in NPCs during IVDD pathogenesis.

### FibroNPCs might be a subpopulation for NP regeneration

Cellular (Cyto) Trajectory Reconstruction Analysis using gene Counts and Expression (CytoTRACE) can accurately uncover the direction of differentiation and predict cell lineage trajectories ^12^. The results indicated the order of the differentiation states as fibroNPCs, adhesion NPCs, effector NPCs, regulatory NPCs, HT-CLNPs, and homeostatic NPCs (Fig. 4A). In addition, we applied the scVelo trajectory algorithms, a novel method developed for recovering the position of each cell in the underlying differentiation processed based on inferred gene-specific rates of transcription, splicing, and degradation. ^38^ The arrows in scVelo indicate the estimated sequence of transcriptomic events (Fig. 4B). The results predict that the fibrosis NPCs may be differentiated into adhesion NPCs and other NPCs subsets. Monocle analysis also revealed that FNPs existed at the start of the pseudospace trajectory. Adhesion NPCs were distributed along the trajectory, and homeostatic NPCs were mainly distributed at the end (Fig. S3A). Together, these findings imply that fibroNPCs possess progenitor properties.

**Figure 4.**
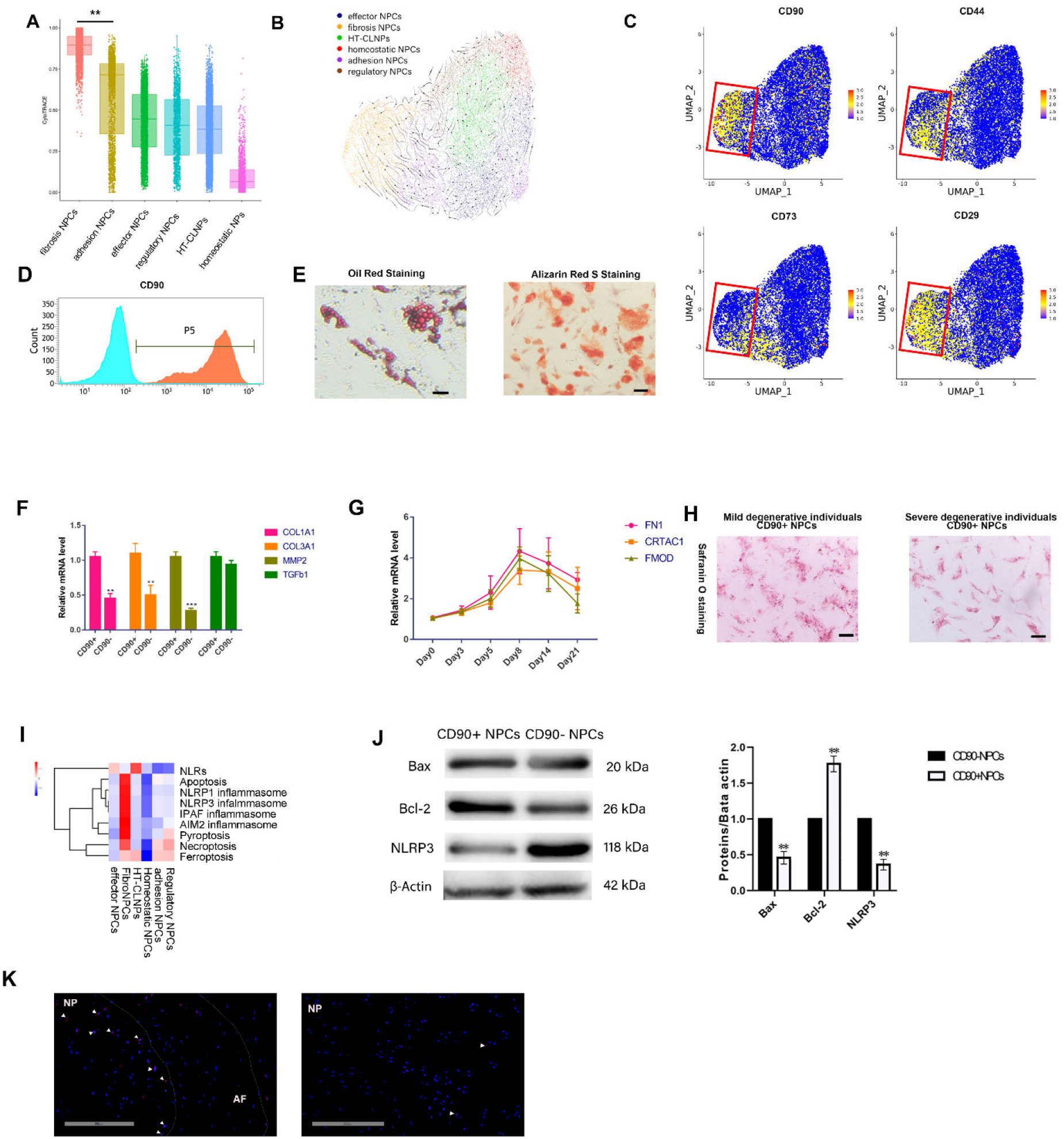
fibroNPCs were identified as the main regeneration subpopulations. **(A)** Plot of the cytoTRACE pseudotime order for the NP subpopulations. The value of cytoTRACE represents the predicted order. **(B)** Visualization for dynamic velocities projected into the UMAP-based embedding. **(C)** The expression of CD90, CD44, CD73, and CD29 in NPCs, the red box represents the region of fibroNPCs on UMAP. **(D)** Histogram to evaluate the relative expression of CD90 after cell sorting. **(E)** Left: oil red staining for CD90+ NPC-induced adipogenic differentiation, respectively (n=3). Scale bar, 100 μm. Right: alizarin red staining for CD90+ NPC-induced osteogenic differentiation (n=3). Scale bar, 100 μm. **(F)** RT-qPCR of fibroNPCs phenotype mRNA levels between CD90+/-NPCs. (n = 3 with mean ± SD shown, *P < 0.05, **P < 0.005.) **(G)** RT-qPCR of adhesion NPCs phenotype at different time points after cultured CD90+NPCs in chondrogenesis induced medium. (n = 3 with mean ± SD shown) **(H)** Safranin O staining of CD90+NPCs from mild (grade I andII) or severely (grade III and IV)degenerative individuals after culturing in chondrogenesis medium for 21 days. **(I)** Correlation of scRNA-seq defined NPC subpopulations with cell death and inflammasome. **(J)** Western blot analysis with representative blots including Bax, Bcl-2m NLRP3 levels in the CD90+/-NPCs. Densitometric analysis is shown as mean ± SD, n =3; *P < 0.05, **P < 0.005. **(K)**Immunofluorescence (IF) visualization of CD90 (red) and nuclei (blue) in degenerative disc tissues induced in rats.

The scVelo analysis also helps to systematically identify putative driver genes as genes characterized by high likelihoods in fibroNPC populations. (Fig.S3B) In other words, these genes may work as candidates for important drivers of the main process in firboNPCs. These genes have been associated with matrix remodeling (Col14A1, col12a1, CHAD, CRTAC1, TNC, Lamb1) ^39^, antioxidation (FTH1) ^40^ and inflammation (S100a8) ^41^.

Since fibroNPCs may have progenitor properties, we analyzed the expression of markers of mesenchymal stem cells (MSCs) in our single-cell data and gated the fibroNPC region with MSCs markers (red box in Fig.4C) on visualized UMAPs. The results show that CD90 was expressed mainly in the fibroNPC region compared to the other MSC markers, CD44, CD73, and CD29. We, therefore, hypothesize that CD90+ NPCs may be the progenitor cells in degenerative NP tissues.

To confirm whether CD90+ NPCs are the progenitor cells in NP, we isolated CD90+NPCs from human degenerated NPCs using microbeads (Fig. 4D). The CD90+NPCs were positive for CD44 and CD29 and negative for CD34 and HLA-DR (Fig. S5A). Oil red staining and Alizarin red staining revealed that CD90+ cells had multipotent capabilities (differentiated into various cell lineages, including osteoblasts and adipocytes) (Fig. 4E). RT-qPCR showed that fibroNPCs phenotype genes (including CLO1A1, COL3A1, MMP2) were significantly elevated in CD90+NPCs compared to CD90-NPCs, however, the expression of TGFb β showed no significant difference between two groups (Fig. 4F). We also cultured CD90+NPCs in the chondrogenesis differentiation medium and found that adhesion NPCs genes, including FN1, CRTAC1, and FMOD, were elevated from day 1 and peaked at day 6 but decreased after that. (Fig. 4G) This result indicates that CD90+NPCs can differentiate into cells with adhesion NPCs phenotypes. Safranin O staining showed a higher chondrogenesis potential for CD90+ NPCs isolated from mild degenerative (grade I and II) individuals compared to those from severe degenerative (grade III and IV)individuals (Fig. 4H). These findings demonstrate that CD90+NPCs expressed phenotypes of fibroNPCs and can serve as progenitor cells in degenerative NP tissues.

Both cell repair and cell death are involved in tissue regeneration ^42^. To explore whether different cell death events occurred in our predictive NPC subsets, we correlated our single-cell data with previous publications ^43, 44^. The correlation analysis showed that fibroNPCs were active for all four cell death types: apoptosis, pyroptosis, necroptosis, and ferroptosis. Inflammasome may be highly involved in cell death in fibroNPCs but not in other cell types. Ferroptosis was evident in HT-CLNPs, adhesion NPCs, and regulatory NPCs. Necroptosis occurred in fibroNPCs, adhesion NPCs, and regulatory NPCs (Fig. 4I). Next. we compared apoptosis and NLRP3-related proteins expressed between CD90+ NPCs and CD90-NPCs; CD90+ cells showed decreased Bax and NLRP3, as well as increased Bcl-2 expression (Fig. 4J). Furthermore, we used immunofluorescence to explore the location of CD90+ NPCs in puncture-induced degenerative rat IVD ^4^. It was found that CD90+ NPCs were enriched in the boundary between NP and AF; only a few positive cells could be detected in the inner area of the discs (Fig. 4K). Therefore, the combined results showing fibroNPCs showed profound cell death activity; however, CD90+NPCs may help NP regeneration in this subpopulation. Overall, fibroNPCs are the major cell population for disc regeneration during IVDD with the persistence of both cell death and progenitors.

### Identification of NP-derived G-MDSCs and validation of their functions

Using the canonical correlation analysis (CCA) method ^45^, we identified neutrophils, G-MDSCs, and G-GMPs based on the expression of hallmark genes (Fig.1D). Apart from previously known cell surface markers for G-MDSCs, ITGAM (CD11b), and OLR1 ^18, 19^, we also identified CD24 as a novel marker. CD11b, OLR1, and CD24 are distinctively highly expressed in G-MDSCs compared to neutrophils and G-GMPs (Fig. 1E). The indicated gene expression in NP tissues at the single-cell level can be found in Fig. 5A. In Fig. 5A, CD45, CD11b, OLR1, and CD24 were not expressed in NPCs, and almost all non-NPCs expressed CD45.

**Figure 5.**
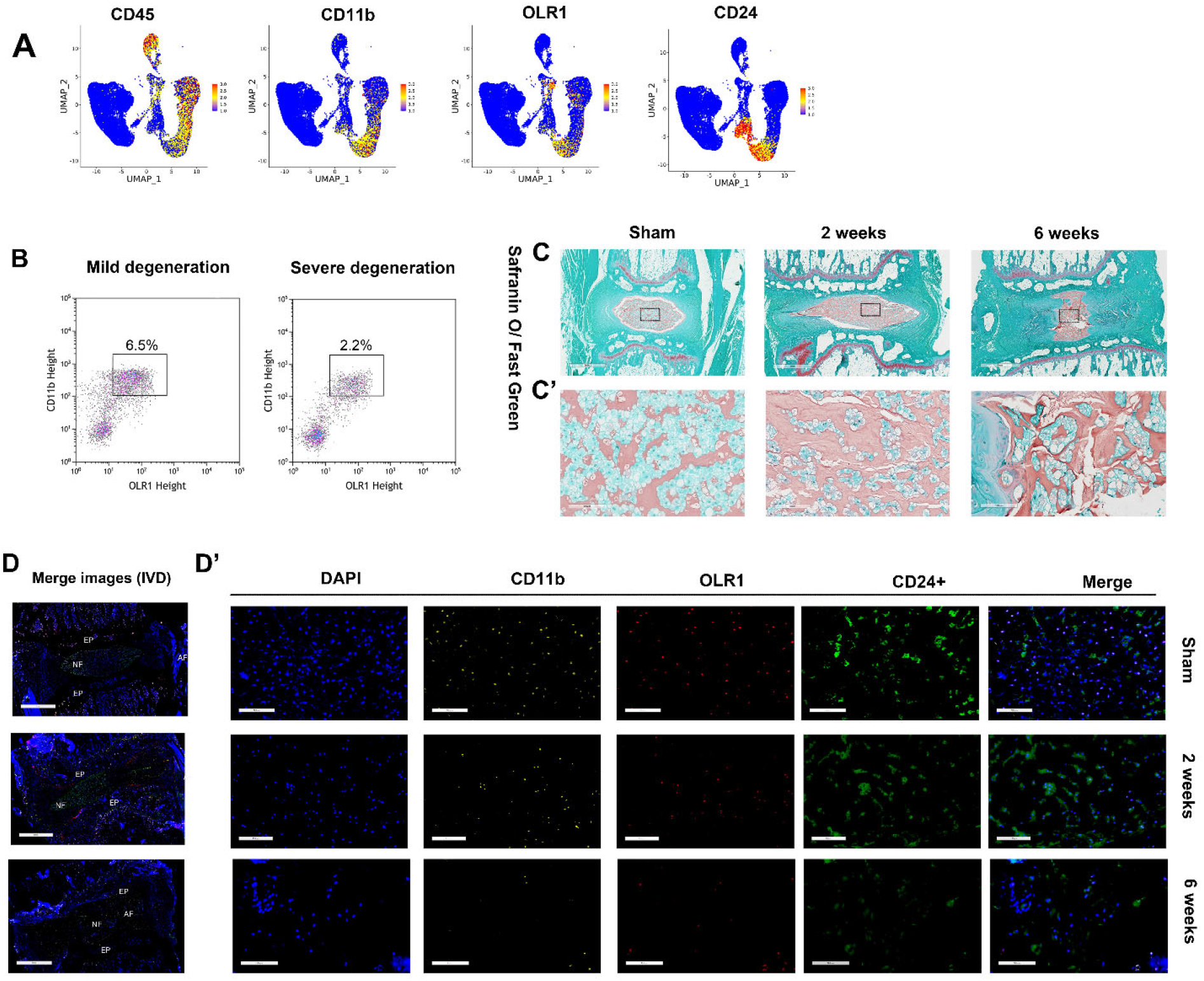
G-MDSCs enriched in mild degenerated NP tissues. **(A)** The expression of CD45, CD11b, OLR1, and CD24 in human NP tissues. **(B)** Quantitative comparison of CD45+CD11b+OLR1+ cells in NP tissues between mild and severe human degenerated discs. **(C)** and **(C’)** shows Safranin O/Fast Green staining of the intervertebral discs sham and experimental rats. Scale bar, 1 mm in C and 100 µm in C’. **(D)** and **(D’)** Merged immunofluorescence staining of DAPI, CD11b, OLR1, and CD24 in the intervertebral discs of sham and experimental rat. Scale bar, 1 mm in D and 100 μm in D’

Our data show that there is an enrichment of G-MDSCs in severely degenerated NP tissues (grade III and IV) compared to mildly degenerative tissues (grade I and II) at the single-cell level. To confirm this finding, we isolated G-MDSCs from human NP tissues via Fluorescence-activated cell sorting (FACS) (Fig. S4). CD45, the marker of hematopoietic cells, CD11b, OLR1, and CD24 were markers used for G-MDSCs sorting. G-MDSCs (CD45+CD11b+OLR1+ cells) decreased by almost three folds in severe degeneration discs (6.5% vs. 2.2%) (Fig. 5B). To further verifyvalidate this finding *in vivo*, we used well-established rat models of IVDD with the needle puncture method ^4^. Histological staining showed notable changes including cell cloning (cell clusters), loss of demarcartiondemarcation between NP and AF, and proteoglycan loss in punctured IVDs, demonstrateddemonstrating progressive IVDD at 2 weeks and 6 weeks (Fig. 5C and C’). Immunofluorescence staining for CD11b, OLR, and CD24 showed the existence of G-MDSCs in rat NPs, and G-MDSCs decreased in the 6-week group relative to the 2-week group (Fig. 5D and D’).

The most typical and important function of G-MDSCs is immunosuppression ^46^. The hallmark of this immunosuppressive activity is the capability of suppressing T cell activation and ROS production ^47, 48^. To functionally validate whether CD45+CD11b+OLR1+CD24+/-cells inhibit immune cell activation, we performed co-cultures with activated T cells (Fig. 6A-B) ^49^. Human NP tissues, both CD45+CD11b+OLR1+CD24+ and CD24-, exhibited T cell suppressive capacity; however, CD24+ cells showed significantly stronger suppressive capacity compared to CD24-cells. (Fig. 6C) We also observed that CD45+CD11b+OLR1+CD24+ cells produced significantly higher amounts of ROS compared to CD24-cells. (Fig. 6D-E) These findings indicate that our identified G-MDSCs (CD45+CD11b+OLR1+ CD24+/-cells) in human NP tissues have T cell suppression and ROS production capabilities.

**Figure 6.**
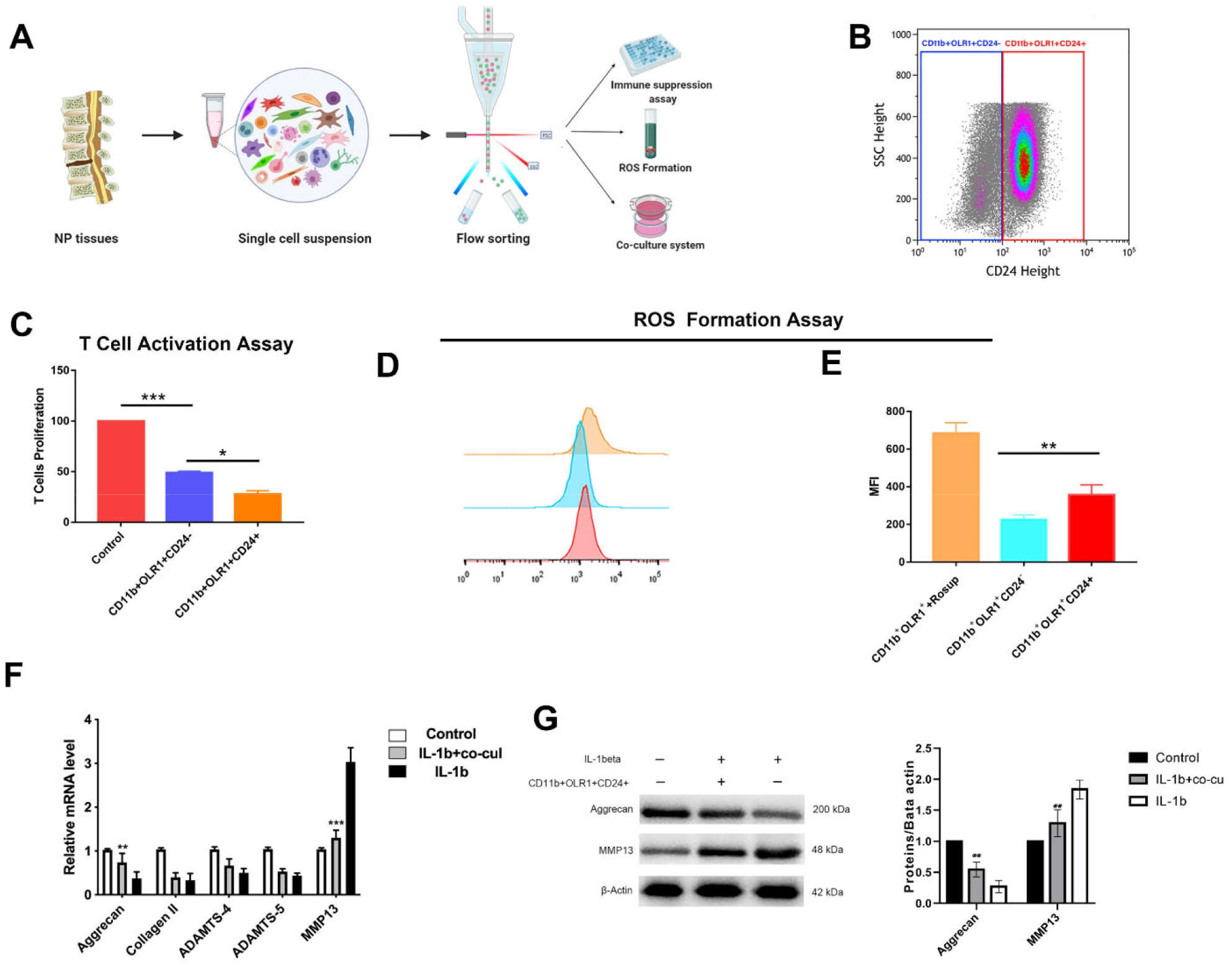
Validation of functions of NP-derived G-MDSCs. **(A)** Schematic workflow of the experimental strategy. **(B)** FACS isolation forCD45+ CD11b+OLR+CD24+ and CD24-cells. **(C)** T cell suppression analysis NP-derived G-MDSC identification. N=3, *P < 0.05, **P < 0.005, ***P<0.001. **(D-E)** CD11b+OLR+CD24+ show increased reactive oxygen species (ROS) formation compared to CD24-cells. Rosup treated cells were used as positive control. N=3, *P < 0.05, **P < 0.005. **(F)** RT-qPCR of degeneration related genes, aggrecan, collagen II, ADAMTS4,5, and MMP13 in untreated, IL-1b+G-MDSCs, and IL-1b alone NPC groups. N=3, *P < 0.05, **P < 0.005, ***P<0.001. **(G)** Western blot analysis with representative blots including aggrecan and MMP13 in untreated, IL-1b+G-MDSCs, and IL-1b alone NPC groups. N=3, *P < 0.05, **P < 0.005, ***P<0.001.

Since G-MDSCs decreased in severely degenerated NP tissues, we hypothesized that G-MDSCs might protect against IVDD. To explore the effects of G-MDSCs on NPCs, G-MDSCs were isolated from mildly degenerated (grade I-II) human NP tissues and co-cultured with non-degenerative NPCs. IL-1β was used to induce NPC degeneration ^50^. Consequently, compared with IL-1β treatment alone, G-MDSCs co-cultured with NPCs showed increased aggrecan expression and significantly lower MMP13 expression (Fig. 6F-G). These findings indicate that G-MDSCs may serve a protective role in NPC degeneration.

### Characterization of cell-to-cell interactions involved in IVDD

Our study showed that NP tissues consist of NPCs and immune cells. We next predicted the cell-cell interaction network among the cell types using CellPhoneDB 2.0. ^51^ Macrophages showed the most interactions with other cell types. Interactions among NPCs were the most active (Fig. 7A).

**Figure 7.**
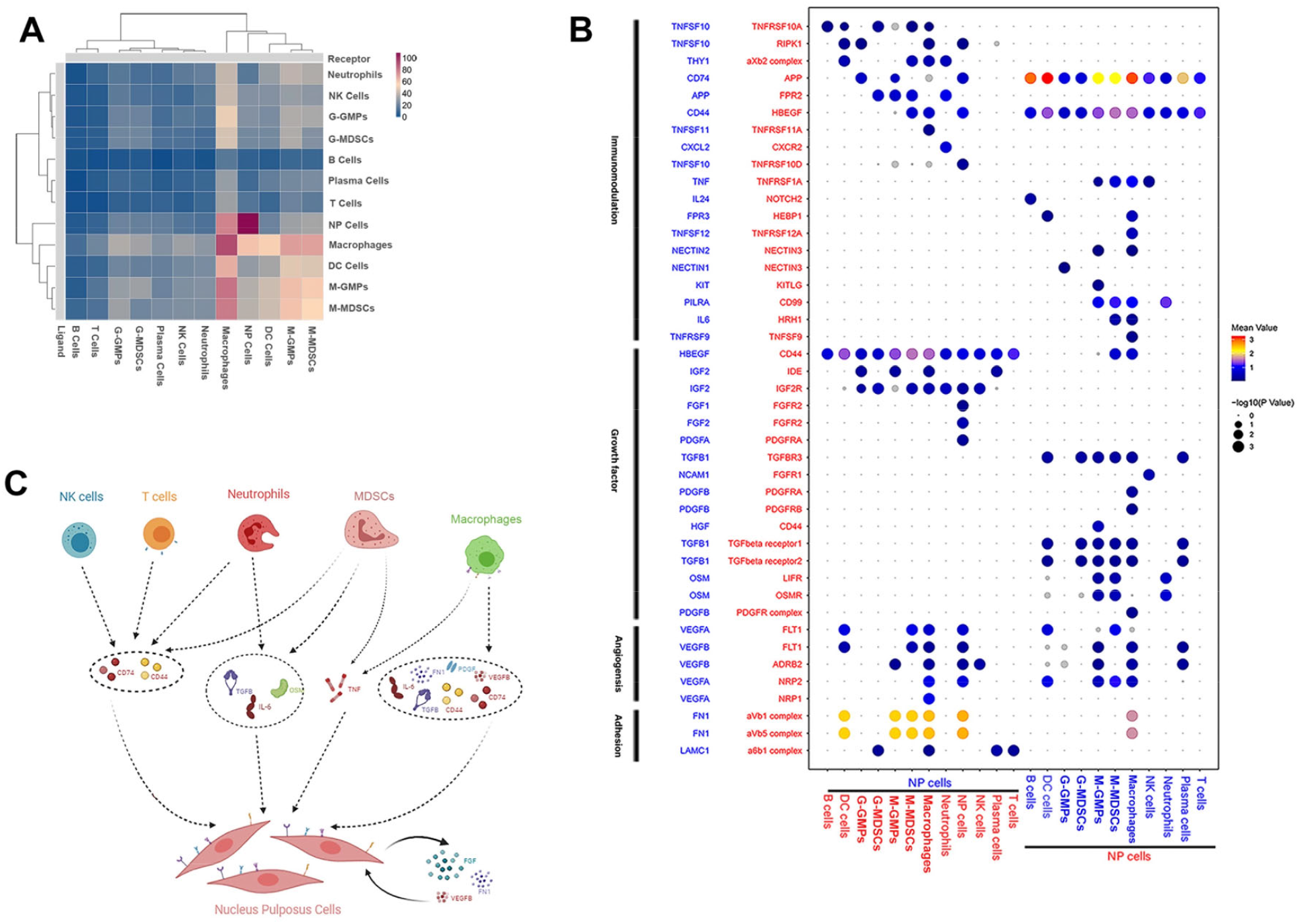
Predicted immune-NPCs regulatory network in IVDD. **(A)** Heatmap showing the number of potential ligand-receptor pairs between cell groups. **(B)** Bubble plots showing ligand-receptor pairs of immunomodulation, growth factors, angiogenesis, and adhesion between NPCs and other cell groups. **(C)** Predicted regulatory network centered on NPCs

CD74/APP and CD44/HBEGF widely existed in the interaction between immune cells and NPCs. NPCs also produced VEGFB and VEGFA, which bind to VEGF receptors (FLT1, ADRB2, NRP1, and NRP2) in both NPCs and immune cells, especially macrophages. This implies that the interaction between macrophages and NPCs is potentially proangiogenic. Notably, a healthy IVD is largely avascular. However, angiogenesis occurs during degeneration ^52^.

It has also been suggested that NPCs produce FN1 and FGF for their receptors. Immunomodulation-related cytokines, such as pro-inflammatory cytokine, TNF, anti-inflammatory cytokines, and IL-6, are expressed by macrophages, neutrophils, and G-MDSCs, and can bind to NPC receptors. Neutrophils and macrophages expressed growth factors, including TGFB, PDGF, and OSM, and showed pro-proliferation effects on NPCs. (Fig. 7B-C). Overall, our results predicted a network between immune cells and degenerative NPCs, which may play a role in inflammation balance, cell proliferation, and angiogenesis in NP. However, the exact mechanisms should be confirmed by further experiments.

## Discussion

Here we provide the first single-cell view of IVDD pathology, profiling NPCs and non-NPCs from NP tissues across individuals with progressive degeneration grades. Notably, we first identified novel cell markers and signatures for verifying each hypothesised NP cluster. Our identified subpopulations showed transcriptomic heterogeneity and potential functional differences at the single-cell level. Effector NPCs were identified and indicated to be active in cellular anabolism and catabolism activities, as they showed a high metabolic rate, including sterol biosynthesis, glycosaminoglycan metabolism, and protein metabolism. During IVDD, the ECM and cells undergo profound metabolic processes ^53^. The QuSAGE analysis indicated effector NPCs are active for “positive regulation of ECM assembly”, which is opposite to fibroNPCs with “negative regulation of ECM assembly”. Thus, effector NPCs may play a role in maintaining ECM homeostasis. It was also suggested that with IVDD progressing, the proportion of effector NPCs decreased, while fibroNPCs proportion increased. Regulatory NPCs are characterized by inflammation and endogenous stimuli responses. The Toll-like signaling pathway, a crucial pathway related to the innate immune system, is active in regulatory NPCs. Moreover, regulatory NPCs have been shown to be one of the main subsets in grade I NP degeneration, suggesting that they may serve as a cell population for immunity maintenance.

During cartilage development, chondrocytes undergo terminal differentiation when they become hypertrophic. It has been widely acknowledged that chondrocyte hypertrophy-like changes play a role in osteoarthritis (OA) ^54^. It has been shown that hypertrophic differentiation occurs in NP tissues during IVDD progression ^55, 56^. Here, we annotated a cell population as “HT-CLNPs” and identified HT-CLNPs-I and HT-CLNPs-II subpopulations possessing different ontologies. The HT-CLNP-I subpopulation was enriched in genes related to negative regulation of apoptosis, response to unfolded protein, and circadian rhythm, while the HT-CLNP-II subpopulation expressed genes involved in ECM organization and disassembly (Table S3). This indicates that HT-CLNP-I may serve as a regulatory cell cluster that may play a protective role in chondrocyte hypertrophy-like events. In support, our data showed decreased HT-CLNP-II proportion in mild degenerative discs (grade I and II). Moreover, a distinct cell marker of HT-CLNP-II, CHRDL2, was found to negatively regulate chondrocyte development by competitively inhibiting BMPs ^57^. BMP/TGFβ is a vital pathway promoting chondrocyte hypertrophy. Here, functional analysis (Fig.4I) showed different cell death events occurring in HT-CLNPs compared to other subpopulations. In HT-CLNPs, ferroptosis, instead of apoptosis, was the main cell death type. HT-CLNPs also showed decreased cell senescence and a low level of SASP. These results revealed potential cell dysfunction events happened in NPCs chondrocyte hypertrophy-like changes.

It has been reported that NP originates from the notochord; however, at about age 4 in humans, notochordal cells disappear, replaced by smaller non-vacuolated cells. More importantly, the origin of these cells is unknown. Previous studies indicated that Tie2 positive (Tie2+) and disialoganglioside 2 positive (GD2+) populations serve as progenitor NPCs ^58, 59^. However, few Tie2+ and GD2+ NPCs can be found in patients aged over 50 ^60^. Interestingly, the single-cell data showed that there are few Tie2+ or GD2+ NPCs in degenerative NP tissues (Fig. S5B). IVDD is an age-related condition; the average age is 59.13+/−12.5 for males and 61.02+/−10.6 for females ^61^. Thus, finding out progenitor cells in aged and degenerative NP tissues is vitally important for understanding tissue homeostasis and regeneration. Our scRNAseq showed that fibroNPCs demonstrated the highest score for progenitor properties and stemness. Increasing evidence shows that apoptotic cells can induce compensatory proliferation and promote regeneration in invertebrates and vertebrates ^42^. The relationship among apoptosis, proliferation, and tissue regeneration has been linked in some animal models, like Drosophila, planarians, newts, and mice ^62, 63^. Cell apoptosis is an important cell death mechanism in IVDD ^37^. Our study showed that apoptosis and inflammasome activity occurred mainly in fibroNPC populations. Moreover, the score for angiogenesis is highest in fibroNPCs and angiogenesis is essential for the growth and regeneration of tissues ^64^. These imply that fibroNPCs are a significant subpopulation for cell regeneration in NP tissues.

We selected CD90+ NPCs as candidate progenitors, which transcriptionally co-express genes for fibroNPCs and MSCs. CD90+NPCs showed decreased apoptosis compared to CD90-NPCs and could differentiate into adipocytes, osteoblasts, and other NPCs atlas (like adhesion NPCs). Furthermore, the CD90+NPCs isolated from mild degeneration individuals (grade I and II) demonstrated increased cell chondrogenesis ability. This study revealed that CD90+ is important for maintaining NP homeostasis and can be used in cell-based scaffolding for NP repair and regeneration. Moreover, CD90+ was found most abundant in the NP/AF boundary regions in rat degenerative disc tissues. Exploring the origin of progenitor cells is of researchers’ interest. Previous descriptive studies found that progenitor and stem cell niche patterns in healthy IVDs are detected in certain AF areas. It was possible that MSCs from the bone marrow niche were involved in NP origination ^65^. Here we hypothesized that a potential niche with CD90+NPCs enriched exists in the NP/AF boundary (Fig. S5C). CD90+NPCs may originate from the AF region, perichondrium region, or from bone marrow-MSCs differentiation and migration. Detailed experiments should be conducted to trace the origin and differentiation of CD90+NPCs in the future.

The IVDs have been identified as immune privilege organs; the steady state of immune privilege is fundamental to organ homeostasis ^66^. This single-cell study showed there are multiple immune cell lines inside the NP, which may play a role in IVDD progression. Based on correlation analyses of previously reported single-cell data, we showed that G-GMPs aligned with bone marrow GMPs, proNeu, and preNeu. G-MDSCs aligned with CXCR2^low^ immNeu and CXCR2^high^ mNeu, and neutrophils aligned with polymorphonuclear leukocytes (PMNs) ^20, 21^. G-MDSC represents a group of immature broadly defined neutrophils with immunosuppressive functions characterized as CD14−CD11b+CD15+/CD66b+ in humans ^67, 68^. MDSCs have been widely studied in cancer, acting as a suppressor of antitumor immune responses ^69^. Accumulation of G-MDSCs has also been reported in cancers and other inflammatory diseases, including inflammatory bowel diseases, rheumatoid arthritis, autoimmune arthritis, and autoimmune hepatitis ^70^. Nevertheless, the involvement of MDSCs in IVDD has not been revealed. MDSCs have a close correlation with neutrophils. Our findings indicated that in NP tissues, G-MDSCs might emerge from GMPs through a differentiation trajectory.

Here, we identified CD24, which can be used in combination with CD11b and OLR1 to detect the presence of G-MDSCs in degenerative NP tissues. These cell surface receptors have been studied for their functions. It should be noted that CD24 was detected to express in NPCs, and CD24+ NPCs decreased with IVDD severity ^71^. Previously, CD24+ NPCs were identified as progenitors in NP^72^. In humans, the percentage of CD24+NPCs decreased significantly with aging or degeneration, which dropped to less than 10% in elderly adults or severely degenerated discs ^73^. In our study, scRNAseq showed that there are few CD24+ NPCs in degenerative discs. That is because there is only one participant below 50 years old in this study. Here, we identified CD24 as a cell marker for detecting NP-derived G-MDSCs when combined with CD11b and OLR1 ^19, 69^. The results of FACS showed that most CD11b+OLR1+ were CD24 positive. Thus, we can conclude that CD24 is mainly expressed in G-MDSCs, but not in NPCs, especially in elderly populations. The immunofluorescence for CD24 in discs of rat models displayed positivity for NPCs, and the percentage of positive cells decreased with IVDD. This might be a result of age and species difference as the rats we employed were 3-month-old. It is currently known that CD24 serves as a costimulatory factor of T cells, regulating their homeostasis and proliferation, and is involved in B cell activation and differentiation ^74, 75^. Our findings showed CD24 can regulate suppression of T cell responses, suggesting CD24 might allow G-MDSCs to directly regulate immune checkpoints in NP tissues. Our single-cell data and animal experiments showed that G-MDSCs substantially decreased in mildly degenerated NP tissues. Our co-culture experiment also suggested G-MDSCs can be a potential treatment option for IVDD.

Although our study provides some previously unrevealed insights into NP tissue biology, there are potential limitations to our study here. Firstly, we need to acknowledge that part of the functional analysis was based on scRNAseq prediction. For instance, the functional difference was among six identified NPC subsets. Other alternative methods should be utilized to estimate the functional biology behind transcriptional changes in future research. Secondly, the dissociation bias when processing tissues for scRNAseq may lead to spurious changes of cell distribution, which can limit our ability to provide an exact numeric description of cell population changes. Several strategies were undertaken to minimize the bias. First, all samples were handled following the same protocol. Second, sample acquisition was carefully made during the surgery to avoid contamination of AF tissues or other tissues. Third, we used multiple methods to show the presence of identified novel cell types, like G-MDSCs. In summary, we have comprehensively decoded the multicellular ecosystem of NP during degeneration. These findings may be leveraged to improve diagnostics and develop preventative strategies for degenerative spinal diseases.

## Acknowledgments

We thank Bo Zhang, Su Wang, Binjin Wang for scientific discussions and assistance with various aspects of the experiments.

## Availability of Data and Materials

All data needed to evaluate the conclusions in the paper are present in the paper and/or the Supplementary Materials. Additional data related to this paper may be requested from the authors. The sequencing data at the NCBI’s Gene expression omnibus (GEO) data repository with the accession ID: GSE165722.

## Supplementary Figures

**Fig. S1.**
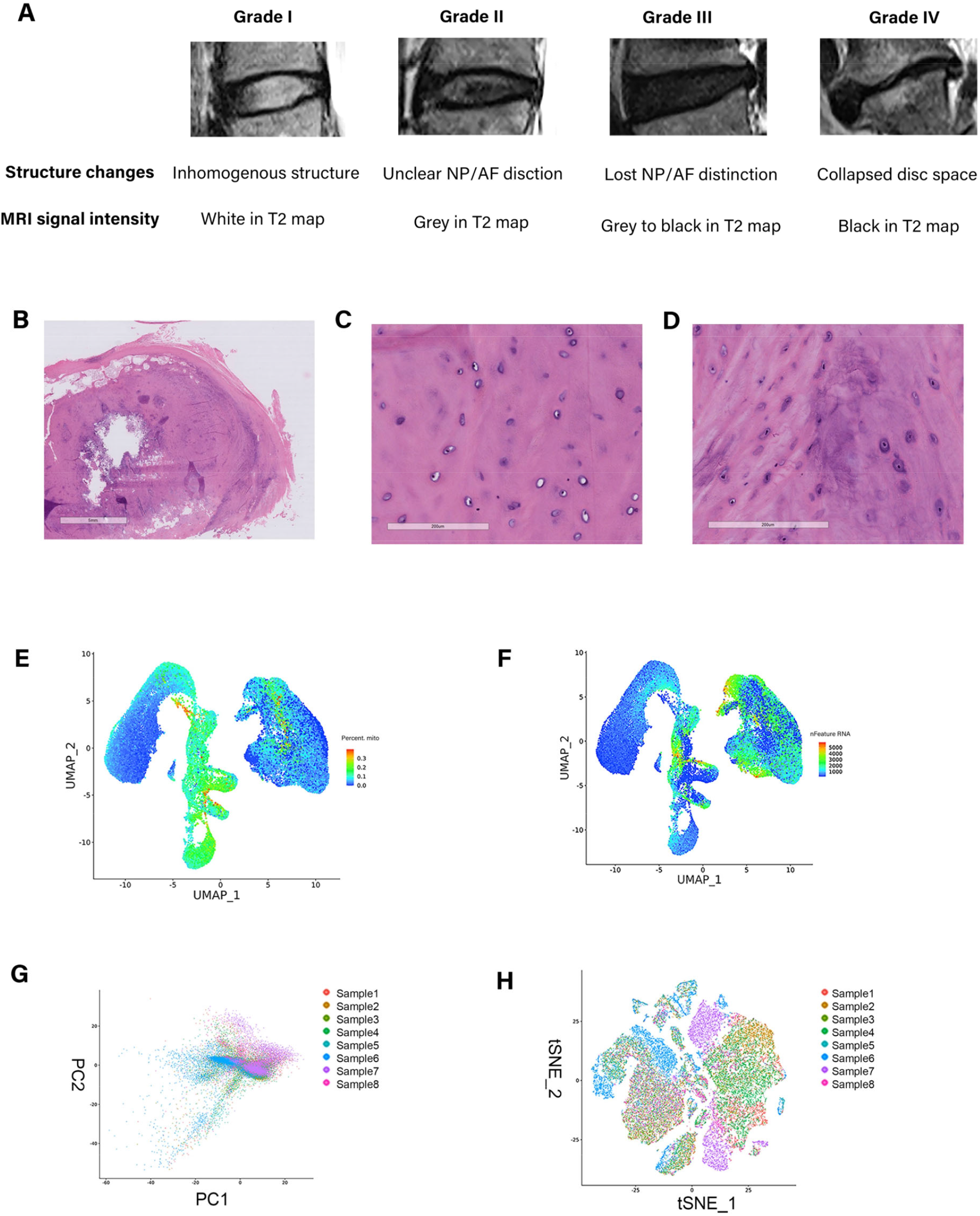
(A) The stage of generation was based on IVD structure and MRI signal intensity. (B-C) Cell quality for single cell sequencing. Hematoxylin and eosin (H&E) staining showed structure of the human disc(B), the NP (C) and the AF(D). Scare bar, 5mm in A; 200um in B and C. (D-H) PCA and tSNE plots of all 39,732 single-cell transcriptomes coloured by individual participants.

**Fig. S2.**
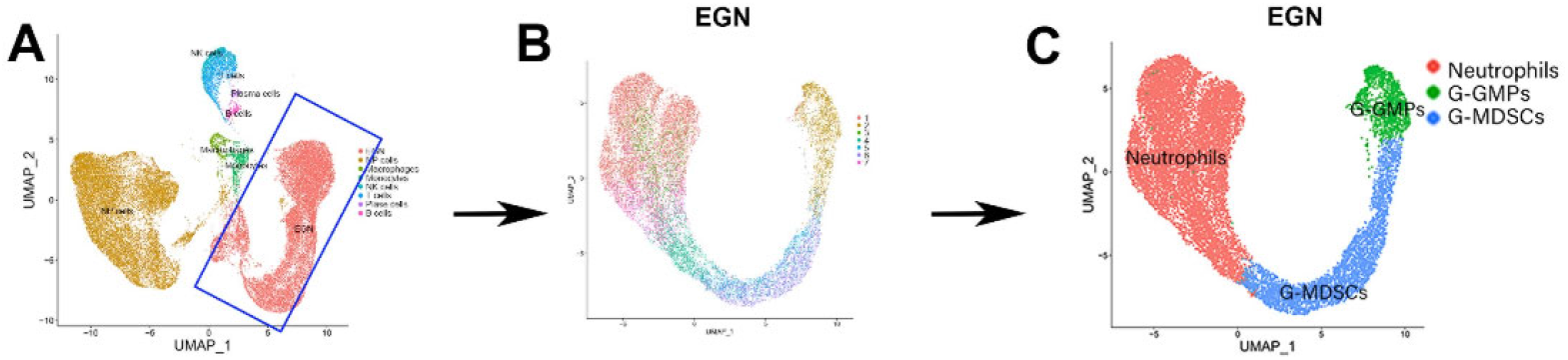
(A) Uniform Manifold Approximation and Projection (UMAP) visualization showing all annotated cells. The blue box is our identified evolutionary group of G-GMPs to neutrophils(EGN). (B) Further clustered and produced 7 transcriptionally distinct pre-clusters from EGN group. (C) Annotated cell neutrophils, G-MDSCs, and G-GMPs based on known gene markers.

**Fig. S3.**
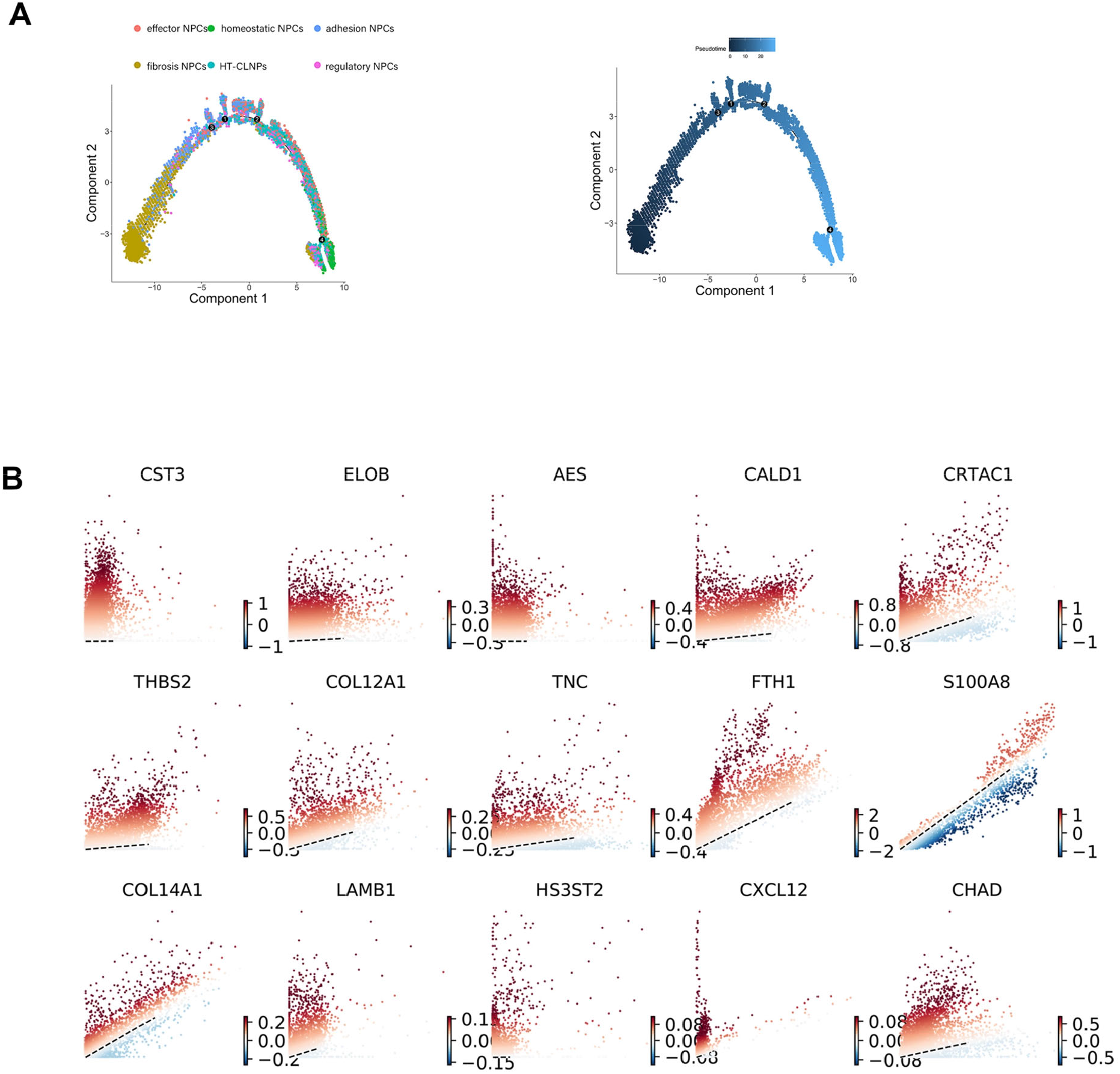
(A) Monocle method reconstruction of pseudospace trajectory for defined NP cell clusters. (B) Identify putative driver genes for fibroNPCs by scVelo.

**Fig. S4.**
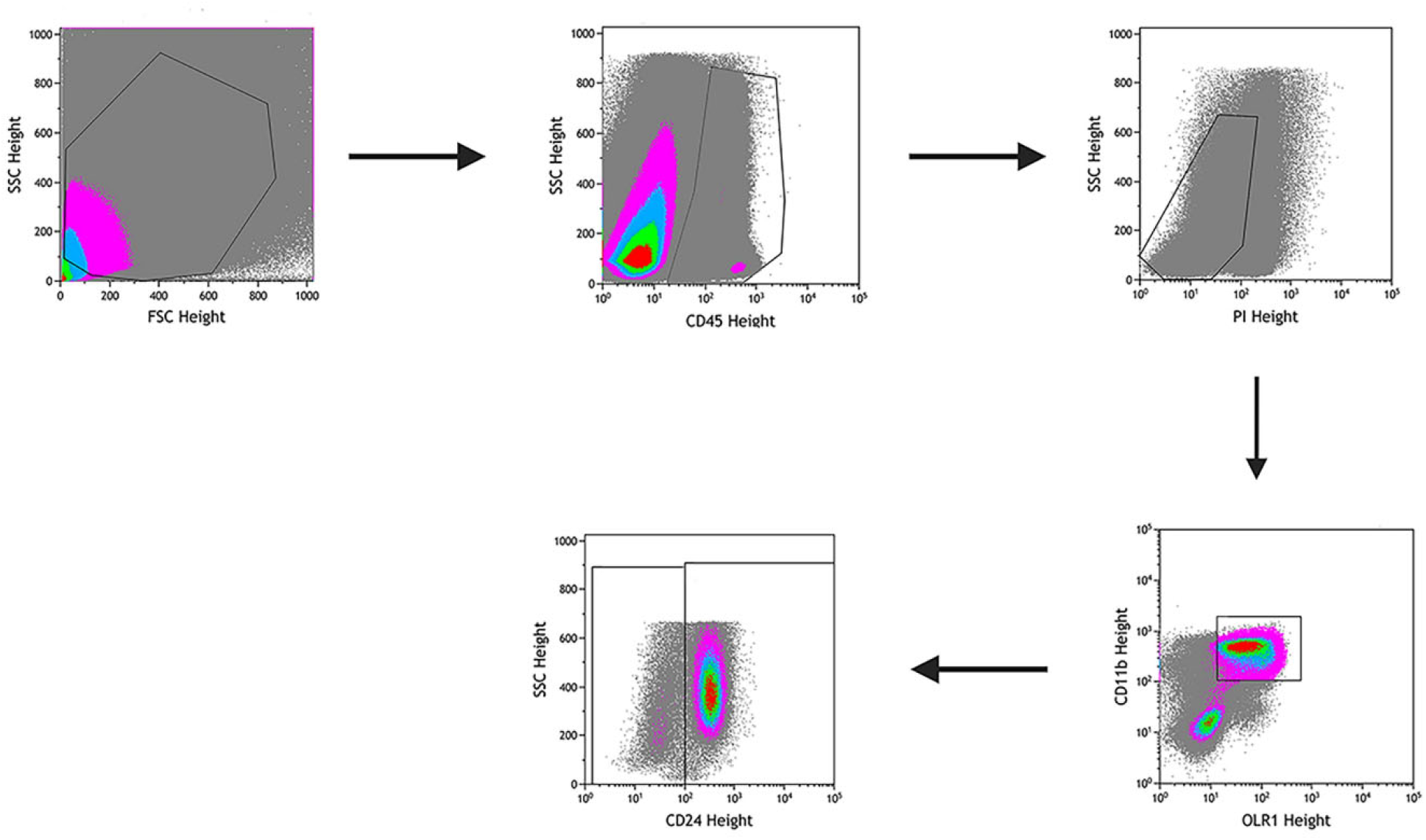
Gating strategies for flow sorting of NP-derived MDSCs.

**Fig. S5.**
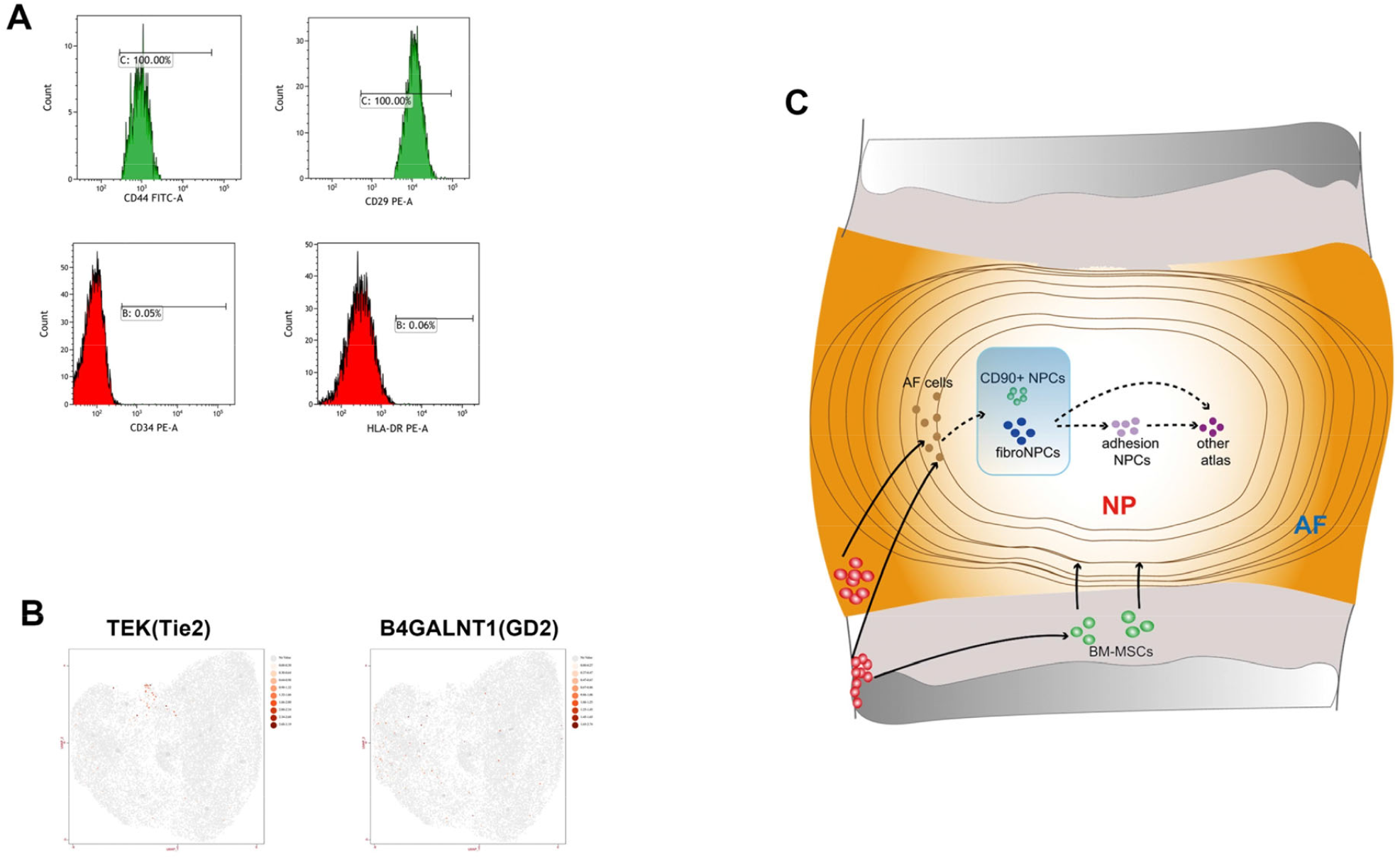
(A) Expression of CD44, CD29, CD34, and HLA-DR in CD90+NPCs. (B)Expression of TEK (Tie2) and Gd2 in human NPCs in UMAP. (B) Schematic graph of the potential hypothetical cellular migration pathways. NP: Nucleus pulposus, AF: Anulus fibrosus.

