## Supplemental Tables for "Single-cell transcriptome profiling reveals multicellular ecosystem of nucleus pulposus during degeneration progression"

Table 1.

| **Patient ID** | **Age (years)** | **Gender** | **Weight**  **(Kg)** | **Reason for surgery** | **Degeneration grade** | **CRP**  **(mg/dL)** | **WBC**  **(109/L)** | **Lymphocyte**  **(109/L)** | **Monocyte (109/L)** |
| --- | --- | --- | --- | --- | --- | --- | --- | --- | --- |
| S1 | 63 | Male | 60 | burst fracture | I | 0.047 | 6.46 | 2.16 | 0.64 |
| S2 | 41 | Male | 73.5 | burst fracture | I | 0.706 | 11.37 | 2.35 | 1.02 |
| S3 | 56 | Female | 62 | Lumbar disc herniation | II | 0.079 | 7.25 | 1.63 | 0.47 |
| S4 | 65 | Female | 76 | Lumbar disc herniation | II | 0.154 | 6.78 | 2.15 | 0.54 |
| S5 | 64 | Female | 50 | Lumbar disc herniation | III | 0.027 | 4.83 | 1.55 | 0.19 |
| S6 | 53 | Female | 60 | Lumbar disc herniation | III | 0.184 | 5.53 | 2.28 | 0.49 |
| S7 | 54 | Male | 68 | Lumbar disc herniation | IV | 0.011 | 10.79 | 3.55 | 0.83 |
| S8 | 56 | Male | 55 | Lumbar disc herniation | IV | 0.022 | 5.75 | 1.86 | 0.61 |

Basic information and characteristics for participants

| **Patient ID** | **Blood glucose (mmol/L)** | **ALT (U/L)** | **AST (U/L)** | **Total Protein (g/L)** | **Serum albumin (g/L)** | **BUN (mmol/L)** | **Cre (μmoI/L)** | **TT (s)** | **APTT (s)** | **INR** |
| --- | --- | --- | --- | --- | --- | --- | --- | --- | --- | --- |
| S1 | 3.77 | 7.4 | 12.7 | 57.3 | 37.4 | 3.3 | 42.4 | 21.6 | 26.7 | 1.01 |
| S2 | 5.40 | 27.7 | 13.4 | 63.3 | 41.0 | 5.0 | 59.1 | 18.4 | 22.8 | 0.92 |
| S3 | 5.05 | 18.7 | 14.8 | 73.3 | 44.9 | 4.8 | 46.4 | 18.1 | 21.9 | 1.02 |
| S4 | 5.41 | 10.6 | 12.3 | 66.5 | 42.1 | 6.1 | 38.0 | 19.8 | 24.4 | 1.07 |
| S5 | 4.52 | 13.6 | 12.6 | 69.0 | 44.0 | 5.5 | 49.1 | 19.5 | 26.2 | 1.00 |
| S6 | 4.66 | 26.5 | 18.3 | 66.3 | 40.7 | 4.6 | 53.8 | 19.7 | 22.4 | 0.99 |
| S7 | 4.84 | 20.6 | 18.0 | 71.5 | 46.3 | 4.9 | 60.3 | 18.2 | 23.6 | 0.96 |
| S8 | 4.70 | 33.7 | 17.0 | 65.8 | 39.4 | 6.4 | 82.0 | 18.6 | 25.9 | 0.92 |

Table 2. DEGs by comparing severe degeneration grade (GradeIII and IV) to mild degeneration grade (Grade I and II)

**Upregulated genes**

| Gene | P value | avg_logFC | pct.1 | pct.2 | p_val_adj |
| --- | --- | --- | --- | --- | --- |
| INHBA | 0 | 1.542242 | 0.787 | 0.491 | 0 |
| FHL2 | 0 | 1.355454 | 0.65 | 0.274 | 0 |
| KLHL21 | 0 | 1.150942 | 0.633 | 0.334 | 0 |
| CYR61 | 0 | 1.149118 | 0.947 | 0.887 | 0 |
| HSPA1B | 0 | 1.059508 | 0.868 | 0.733 | 0 |
| NFATC2 | 0 | 1.027662 | 0.497 | 0.173 | 0 |
| HSPH1 | 0 | 0.983483 | 0.813 | 0.634 | 0 |
| CTGF | 0 | 0.979719 | 0.957 | 0.877 | 0 |
| MMP13 | 0 | 0.948365 | 0.822 | 0.626 | 0 |
| HSP90AA1 | 0 | 0.899521 | 0.964 | 0.896 | 0 |
| RTN4 | 0 | 0.759251 | 0.836 | 0.694 | 0 |
| HSPA8 | 0 | 0.650494 | 0.882 | 0.8 | 0 |
| ACTG1 | 0 | 0.627876 | 0.979 | 0.96 | 0 |
| ANXA5 | 0 | 0.608453 | 0.908 | 0.839 | 0 |
| LUM | 0 | 0.58595 | 0.984 | 0.97 | 0 |
| H3F3B | 0 | 0.496407 | 0.963 | 0.94 | 0 |
| HSPA1A | 5.2E-284 | 0.735349 | 0.91 | 0.866 | 1.1E-279 |
| GFPT2 | 6.9E-261 | 0.734121 | 0.682 | 0.489 | 1.5E-256 |
| KLF4 | 7E-258 | 0.702163 | 0.905 | 0.846 | 1.5E-253 |
| ATP1B1 | 6.7E-252 | 0.710599 | 0.614 | 0.392 | 1.5E-247 |
| PCOLCE2 | 1.2E-250 | 0.563587 | 0.869 | 0.752 | 2.6E-246 |
| EMP1 | 7.9E-244 | 0.770794 | 0.907 | 0.84 | 1.7E-239 |
| SLC39A14 | 9.2E-241 | 0.606197 | 0.801 | 0.68 | 2E-236 |
| HERPUD1 | 1.3E-237 | 0.515521 | 0.873 | 0.801 | 2.8E-233 |
| PLOD2 | 5.6E-227 | 0.471544 | 0.866 | 0.805 | 1.2E-222 |
| HSPB8 | 1.8E-224 | 0.780165 | 0.687 | 0.502 | 4E-220 |
| ZSWIM6 | 1.8E-219 | 0.548961 | 0.309 | 0.088 | 3.8E-215 |
| BTF3 | 3.4E-219 | 0.457349 | 0.863 | 0.785 | 7.4E-215 |
| ANXA1 | 2.2E-216 | 0.58282 | 0.928 | 0.894 | 4.7E-212 |
| ACKR3 | 4.6E-215 | 0.909398 | 0.595 | 0.397 | 1E-210 |
| MYADM | 1.2E-214 | 0.682027 | 0.739 | 0.614 | 2.5E-210 |
| ACTB | 1.6E-214 | 0.357681 | 0.965 | 0.945 | 3.4E-210 |
| AMOTL2 | 1.1E-212 | 0.883341 | 0.388 | 0.169 | 2.5E-208 |
| FOXC2 | 1.9E-210 | 0.758533 | 0.43 | 0.208 | 4.2E-206 |
| TXN | 1.2E-201 | 0.630013 | 0.708 | 0.571 | 2.6E-197 |
| UAP1 | 7.7E-201 | 0.602079 | 0.708 | 0.556 | 1.7E-196 |
| TM4SF1 | 4.7E-200 | 0.936974 | 0.488 | 0.282 | 1E-195 |
| RYBP | 1.2E-199 | 0.648708 | 0.654 | 0.479 | 2.7E-195 |
| FMOD | 1.1E-195 | 0.535043 | 0.908 | 0.89 | 2.3E-191 |
| RPL30 | 1.8E-186 | 0.341223 | 0.949 | 0.941 | 3.9E-182 |
| GPRC5A | 7.1E-186 | 0.670452 | 0.596 | 0.403 | 1.5E-181 |
| HSPD1 | 1.2E-184 | 0.574874 | 0.775 | 0.666 | 2.7E-180 |
| FN1 | 4.3E-180 | 0.740568 | 0.968 | 0.95 | 9.4E-176 |
| TMED2 | 2.4E-177 | 0.583225 | 0.672 | 0.564 | 5.2E-173 |
| RAN | 1.5E-174 | 0.462583 | 0.799 | 0.715 | 3.3E-170 |
| SERPINE2 | 2E-174 | 0.721224 | 0.845 | 0.773 | 4.4E-170 |
| CTNNB1 | 1.9E-167 | 0.511848 | 0.66 | 0.526 | 4.2E-163 |
| IL11 | 7.6E-165 | 0.928692 | 0.287 | 0.1 | 1.6E-160 |
| OAT | 1E-163 | 0.473536 | 0.728 | 0.608 | 2.3E-159 |
| CD55 | 6.3E-161 | 0.717146 | 0.707 | 0.576 | 1.4E-156 |

**Downregulated genes**

| Gene | P value | avg_logFC | pct.1 | pct.2 | p_val_adj |
| --- | --- | --- | --- | --- | --- |
| CCL4 | 0 | -0.28029 | 0.019 | 0.2 | 0 |
| CCL3 | 0 | -0.28546 | 0.022 | 0.194 | 0 |
| BARX1 | 0 | -0.28722 | 0.004 | 0.148 | 0 |
| CCL3L3 | 0 | -0.30709 | 0.026 | 0.22 | 0 |
| SLC25A1 | 0 | -0.32756 | 0.009 | 0.169 | 0 |
| C19orf70 | 0 | -0.35763 | 0.008 | 0.172 | 0 |
| EIF3G | 0 | -0.36931 | 0.014 | 0.211 | 0 |
| AIP | 0 | -0.38403 | 0.009 | 0.196 | 0 |
| GNB2 | 0 | -0.38455 | 0.029 | 0.241 | 0 |
| NT5DC2 | 0 | -0.38716 | 0.03 | 0.228 | 0 |
| ASS1 | 0 | -0.39016 | 0.047 | 0.25 | 0 |
| RFNG | 0 | -0.39021 | 0.052 | 0.263 | 0 |
| LAMTOR4 | 0 | -0.40186 | 0.059 | 0.274 | 0 |
| CCL4L2 | 0 | -0.40452 | 0.017 | 0.249 | 0 |
| SORBS3 | 0 | -0.40863 | 0.042 | 0.261 | 0 |
| ARL6IP4 | 0 | -0.41084 | 0.031 | 0.241 | 0 |
| HLA-DRA | 0 | -0.42087 | 0.016 | 0.26 | 0 |
| NRBP1 | 0 | -0.42194 | 0.048 | 0.277 | 0 |
| HLA-DRB1 | 0 | -0.44289 | 0.012 | 0.249 | 0 |
| YIF1A | 0 | -0.45561 | 0.053 | 0.286 | 0 |
| SH3BGRL3 | 0 | -0.46366 | 0.044 | 0.258 | 0 |
| ATP6V0B | 0 | -0.46735 | 0.091 | 0.334 | 0 |
| GUK1 | 0 | -0.48434 | 0.049 | 0.281 | 0 |
| ANAPC11 | 0 | -0.49102 | 0.087 | 0.323 | 0 |
| H2AFJ | 0 | -0.49279 | 0.034 | 0.256 | 0 |
| COX5B | 0 | -0.49904 | 0.077 | 0.305 | 0 |
| GAS6 | 0 | -0.5001 | 0.028 | 0.244 | 0 |
| MYL9 | 0 | -0.50117 | 0.029 | 0.238 | 0 |
| BSG | 0 | -0.50177 | 0.095 | 0.37 | 0 |
| METTL26 | 0 | -0.50619 | 0.028 | 0.274 | 0 |
| LY6E | 0 | -0.50741 | 0.039 | 0.281 | 0 |
| ADIRF | 0 | -0.5311 | 0.03 | 0.264 | 0 |
| MMP24OS | 0 | -0.54508 | 0.042 | 0.304 | 0 |
| TUBB4B | 0 | -0.5488 | 0.112 | 0.379 | 0 |
| LAMB2 | 0 | -0.55969 | 0.095 | 0.35 | 0 |
| NOP53 | 0 | -0.56125 | 0.119 | 0.408 | 0 |
| PHPT1 | 0 | -0.56333 | 0.013 | 0.281 | 0 |
| RCN3 | 0 | -0.56512 | 0.164 | 0.433 | 0 |
| RPS5 | 0 | -0.56633 | 0.817 | 0.898 | 0 |
| GPX1 | 0 | -0.57485 | 0.108 | 0.38 | 0 |
| NUCB1 | 0 | -0.58197 | 0.217 | 0.506 | 0 |
| FXYD1 | 0 | -0.59014 | 0.033 | 0.28 | 0 |
| PRDX2 | 0 | -0.59675 | 0.076 | 0.353 | 0 |
| RPL3 | 0 | -0.60441 | 0.242 | 0.558 | 0 |
| EIF5A | 0 | -0.60456 | 0.076 | 0.392 | 0 |
| FXYD5 | 0 | -0.61747 | 0.175 | 0.461 | 0 |
| ID3 | 0 | -0.62269 | 0.177 | 0.488 | 0 |
| LTBP3 | 0 | -0.62545 | 0.088 | 0.376 | 0 |
| MDFI | 0 | -0.63195 | 0.035 | 0.267 | 0 |
| ALDOA | 0 | -0.63467 | 0.489 | 0.722 | 0 |
| RPL13 | 0 | -0.63534 | 0.775 | 0.892 | 0 |

**Table 3 GO Terms for NP subpopulations**

**ENPs GO terms**

| **GOID** | **GOTerm** | **P-Value** |
| --- | --- | --- |
| GO:0016126 | sterol biosynthetic process | 2.432E-11 |
| GO:0006695 | cholesterol biosynthetic process | 5.795E-10 |
| GO:0030198 | extracellular matrix organization | 3.136E-08 |
| GO:0044281 | small molecule metabolic process | 4.528E-08 |
| GO:0001558 | regulation of cell growth | 2.285E-06 |
| GO:0001501 | skeletal system development | 3.133E-06 |
| GO:0006694 | steroid biosynthetic process | 5.308E-06 |
| GO:0008203 | cholesterol metabolic process | 9.283E-06 |
| GO:0008202 | steroid metabolic process | 1.061E-05 |
| GO:0006048 | UDP-N-acetylglucosamine biosynthetic process | 1.904E-05 |
| GO:0008299 | isoprenoid biosynthetic process | 2.17E-05 |
| GO:0010628 | positive regulation of gene expression | 3.007E-05 |
| GO:0008285 | negative regulation of cell proliferation | 3.96E-05 |
| GO:0009405 | pathogenesis | 5.295E-05 |
| GO:0072593 | reactive oxygen species metabolic process | 6.08E-05 |
| GO:0006629 | lipid metabolic process | 9.184E-05 |
| GO:0048661 | positive regulation of smooth muscle cell proliferation | 9.297E-05 |
| GO:0033173 | calcineurin-NFAT signaling cascade | 0.0001037 |
| GO:0030199 | collagen fibril organization | 0.0001053 |
| GO:0042127 | regulation of cell proliferation | 0.0001366 |
| GO:0001525 | angiogenesis | 0.0001439 |
| GO:0014070 | response to organic cyclic compound | 0.0001497 |
| GO:0010955 | negative regulation of protein processing | 0.0001541 |
| GO:0042340 | keratan sulfate catabolic process | 0.0001541 |
| GO:0006011 | UDP-glucose metabolic process | 0.0001563 |
| GO:0007568 | aging | 0.0001871 |
| GO:0006633 | fatty acid biosynthetic process | 0.0002058 |
| GO:0043434 | response to peptide hormone | 0.000223 |
| GO:0055114 | oxidation-reduction process | 0.0002321 |
| GO:0005975 | carbohydrate metabolic process | 0.0003237 |
| GO:0043065 | positive regulation of apoptotic process | 0.0003622 |
| GO:0030203 | glycosaminoglycan metabolic process | 0.0003728 |
| GO:0043066 | negative regulation of apoptotic process | 0.000445 |
| GO:0003331 | positive regulation of extracellular matrix constituent secretion | 0.0004652 |
| GO:0006065 | UDP-glucuronate biosynthetic process | 0.0004652 |
| GO:2000544 | regulation of endothelial cell chemotaxis to fibroblast growth factor | 0.0004652 |
| GO:0042542 | response to hydrogen peroxide | 0.0004732 |
| GO:0032270 | positive regulation of cellular protein metabolic process | 0.0005055 |
| GO:0006915 | apoptotic process | 0.0006357 |
| GO:0010035 | response to inorganic substance | 0.0006375 |
| GO:0008284 | positive regulation of cell proliferation | 0.0006802 |
| GO:0044255 | cellular lipid metabolic process | 0.0007222 |
| GO:0060591 | chondroblast differentiation | 0.0009226 |
| GO:1903169 | regulation of calcium ion transmembrane transport | 0.0009226 |
| GO:0006933 | negative regulation of cell adhesion involved in substrate-bound cell migration | 0.0009226 |
| GO:0051387 | negative regulation of neurotrophin TRK receptor signaling pathway | 0.0009226 |
| GO:0009314 | response to radiation | 0.0009242 |
| GO:0060291 | long-term synaptic potentiation | 0.0010297 |
| GO:0045444 | fat cell differentiation | 0.0013192 |
| GO:0031394 | positive regulation of prostaglandin biosynthetic process | 0.001525 |

**FNPs GO Terms**

| **GOID** | **GOTerm** | **P-Value** |
| --- | --- | --- |
| GO:0030198 | extracellular matrix organization | 8.663E-42 |
| GO:0007155 | cell adhesion | 9.642E-28 |
| GO:0022617 | extracellular matrix disassembly | 1.776E-23 |
| GO:0030574 | collagen catabolic process | 5.312E-19 |
| GO:0007411 | axon guidance | 2.226E-16 |
| GO:0030199 | collagen fibril organization | 8.14E-16 |
| GO:0001525 | angiogenesis | 1.304E-12 |
| GO:0002576 | platelet degranulation | 1.732E-11 |
| GO:0030168 | platelet activation | 9.024E-11 |
| GO:0007507 | heart development | 1.504E-10 |
| GO:0006928 | cellular component movement | 2.848E-10 |
| GO:0007596 | blood coagulation | 2.876E-09 |
| GO:0008285 | negative regulation of cell proliferation | 7.397E-09 |
| GO:0016477 | cell migration | 3.908E-07 |
| GO:0001558 | regulation of cell growth | 4.529E-07 |
| GO:0009612 | response to mechanical stimulus | 6.42E-07 |
| GO:0048010 | vascular endothelial growth factor receptor signaling pathway | 1.441E-06 |
| GO:0051017 | actin filament bundle assembly | 1.628E-06 |
| GO:0001649 | osteoblast differentiation | 2.019E-06 |
| GO:0001666 | response to hypoxia | 2.039E-06 |
| GO:0008283 | cell proliferation | 2.174E-06 |
| GO:0035987 | endodermal cell differentiation | 3.631E-06 |
| GO:0007162 | negative regulation of cell adhesion | 3.942E-06 |
| GO:0007160 | cell-matrix adhesion | 4.348E-06 |
| GO:0030336 | negative regulation of cell migration | 4.63E-06 |
| GO:0051764 | actin crosslink formation | 5.041E-06 |
| GO:0071230 | cellular response to amino acid stimulus | 5.525E-06 |
| GO:0050900 | leukocyte migration | 6.418E-06 |
| GO:0048013 | ephrin receptor signaling pathway | 7.117E-06 |
| GO:0030335 | positive regulation of cell migration | 7.697E-06 |
| GO:0001974 | blood vessel remodeling | 8.644E-06 |
| GO:0016525 | negative regulation of angiogenesis | 9E-06 |
| GO:0001501 | skeletal system development | 9.137E-06 |
| GO:0071305 | cellular response to vitamin D | 1.097E-05 |
| GO:0010811 | positive regulation of cell-substrate adhesion | 1.102E-05 |
| GO:0045669 | positive regulation of osteoblast differentiation | 1.291E-05 |
| GO:0048813 | dendrite morphogenesis | 1.394E-05 |
| GO:0051493 | regulation of cytoskeleton organization | 1.57E-05 |
| GO:0042060 | wound healing | 1.585E-05 |
| GO:0009611 | response to wounding | 1.821E-05 |
| GO:0006936 | muscle contraction | 2.038E-05 |
| GO:0006987 | activation of signaling protein activity involved in unfolded protein response | 2.151E-05 |
| GO:0043491 | protein kinase B signaling | 2.758E-05 |
| GO:0046718 | viral entry into host cell | 3.305E-05 |
| GO:0001938 | positive regulation of endothelial cell proliferation | 3.468E-05 |
| GO:0032964 | collagen biosynthetic process | 3.531E-05 |
| GO:0060674 | placenta blood vessel development | 3.762E-05 |
| GO:0031532 | actin cytoskeleton reorganization | 5.90377E-05 |
| GO:0006898 | receptor-mediated endocytosis | 6.00706E-05 |
| GO:0030036 | actin cytoskeleton organization | 6.00706E-05 |

**HomNPs GO Terms**

| **GOID** | **GOTerm** | **P-Value** |
| --- | --- | --- |
| GO:0006614 | SRP-dependent cotranslational protein targeting to membrane | 2.963E-89 |
| GO:0006415 | translational termination | 4.999E-86 |
| GO:0006414 | translational elongation | 7.153E-80 |
| GO:0019083 | viral transcription | 1.067E-76 |
| GO:0006412 | translation | 1.077E-76 |
| GO:0000184 | nuclear-transcribed mRNA catabolic process, nonsense-mediated decay | 5.322E-75 |
| GO:0019058 | viral life cycle | 9.812E-75 |
| GO:0006413 | translational initiation | 1.227E-74 |
| GO:0016070 | RNA metabolic process | 1.437E-71 |
| GO:0016071 | mRNA metabolic process | 3.898E-67 |
| GO:0010467 | gene expression | 9.55E-63 |
| GO:0016032 | viral process | 4.056E-53 |
| GO:0044267 | cellular protein metabolic process | 3.626E-47 |
| GO:0022904 | respiratory electron transport chain | 2.26E-21 |
| GO:0044237 | cellular metabolic process | 1.722E-18 |
| GO:0002181 | cytoplasmic translation | 5.217E-15 |
| GO:0042274 | ribosomal small subunit biogenesis | 1.254E-11 |
| GO:0000027 | ribosomal large subunit assembly | 1.824E-11 |
| GO:0000028 | ribosomal small subunit assembly | 2.147E-10 |
| GO:0002479 | antigen processing and presentation of exogenous peptide antigen via MHC class I, TAP-dependent | 1.431E-09 |
| GO:0042590 | antigen processing and presentation of exogenous peptide antigen via MHC class I | 2.655E-09 |
| GO:0006120 | mitochondrial electron transport, NADH to ubiquinone | 1.106E-08 |
| GO:0006364 | rRNA processing | 1.348E-08 |
| GO:1902600 | hydrogen ion transmembrane transport | 1.963E-08 |
| GO:0006367 | transcription initiation from RNA polymerase II promoter | 2.211E-08 |
| GO:0051437 | positive regulation of ubiquitin-protein ligase activity involved in mitotic cell cycle | 2.365E-08 |
| GO:0051439 | regulation of ubiquitin-protein ligase activity involved in mitotic cell cycle | 4.824E-08 |
| GO:0000398 | mRNA splicing, via spliceosome | 5.565E-08 |
| GO:0051436 | negative regulation of ubiquitin-protein ligase activity involved in mitotic cell cycle | 9.745E-08 |
| GO:0038061 | NIK/NF-kappaB signaling | 1.169E-07 |
| GO:0031145 | anaphase-promoting complex-dependent proteasomal ubiquitin-dependent protein catabolic process | 1.282E-07 |
| GO:0002474 | antigen processing and presentation of peptide antigen via MHC class I | 2.331E-07 |
| GO:0006977 | DNA damage response, signal transduction by p53 class mediator resulting in cell cycle arrest | 8.067E-07 |
| GO:0033209 | tumor necrosis factor-mediated signaling pathway | 9.747E-07 |
| GO:0000387 | spliceosomal snRNP assembly | 2.742E-06 |
| GO:0034660 | ncRNA metabolic process | 4.53E-06 |
| GO:0006521 | regulation of cellular amino acid metabolic process | 5.739E-06 |
| GO:0050434 | positive regulation of viral transcription | 6.873E-06 |
| GO:0042776 | mitochondrial ATP synthesis coupled proton transport | 1.66E-05 |
| GO:0044281 | small molecule metabolic process | 1.913E-05 |
| GO:0006446 | regulation of translational initiation | 2.391E-05 |
| GO:0000082 | G1/S transition of mitotic cell cycle | 2.8E-05 |
| GO:0010499 | proteasomal ubiquitin-independent protein catabolic process | 3.292E-05 |
| GO:0019068 | virion assembly | 3.292E-05 |
| GO:0000209 | protein polyubiquitination | 3.844E-05 |
| GO:0042254 | ribosome biogenesis | 4.293E-05 |
| GO:0001731 | formation of translation preinitiation complex | 4.472E-05 |
| GO:0006368 | transcription elongation from RNA polymerase II promoter | 4.798E-05 |
| GO:0021888 | hypothalamus gonadotrophin-releasing hormone neuron development | 4.944E-05 |
| GO:0002223 | stimulatory C-type lectin receptor signaling pathway | 5.147E-05 |

**HT-CLNPs GO Terms**

| **GOID** | **GOTerm** | **P-Value** |
| --- | --- | --- |
| GO:0043066 | negative regulation of apoptotic process | 9.969E-11 |
| GO:0044267 | cellular protein metabolic process | 1.093E-09 |
| GO:0000122 | negative regulation of transcription from RNA polymerase II promoter | 1.504E-09 |
| GO:0006413 | translational initiation | 2.324E-09 |
| GO:0045944 | positive regulation of transcription from RNA polymerase II promoter | 2.996E-09 |
| GO:0006366 | transcription from RNA polymerase II promoter | 4.917E-09 |
| GO:0010467 | gene expression | 1.639E-07 |
| GO:0007623 | circadian rhythm | 3.873E-07 |
| GO:0001501 | skeletal system development | 4.605E-07 |
| GO:1901653 | cellular response to peptide | 8.175E-07 |
| GO:0043065 | positive regulation of apoptotic process | 1.082E-06 |
| GO:0007179 | transforming growth factor beta receptor signaling pathway | 4.329E-06 |
| GO:0009409 | response to cold | 6.217E-06 |
| GO:0006986 | response to unfolded protein | 6.218E-06 |
| GO:0006446 | regulation of translational initiation | 7.305E-06 |
| GO:0006412 | translation | 8.13E-06 |
| GO:0016071 | mRNA metabolic process | 8.787E-06 |
| GO:0009612 | response to mechanical stimulus | 1.109E-05 |
| GO:0032496 | response to lipopolysaccharide | 1.285E-05 |
| GO:0008284 | positive regulation of cell proliferation | 2.057E-05 |
| GO:0035914 | skeletal muscle cell differentiation | 2.28E-05 |
| GO:0071479 | cellular response to ionizing radiation | 2.342E-05 |
| GO:0014070 | response to organic cyclic compound | 2.375E-05 |
| GO:0030968 | endoplasmic reticulum unfolded protein response | 2.616E-05 |
| GO:0070301 | cellular response to hydrogen peroxide | 4.16E-05 |
| GO:0006415 | translational termination | 5.61E-05 |
| GO:0006414 | translational elongation | 6.171E-05 |
| GO:0006355 | regulation of transcription, DNA-templated | 7.051E-05 |
| GO:0006351 | transcription, DNA-templated | 7.28E-05 |
| GO:0006457 | protein folding | 7.913E-05 |
| GO:0051726 | regulation of cell cycle | 8.277E-05 |
| GO:0042493 | response to drug | 8.474E-05 |
| GO:0010501 | RNA secondary structure unwinding | 9.475E-05 |
| GO:0006915 | apoptotic process | 9.491E-05 |
| GO:0006950 | response to stress | 9.576E-05 |
| GO:0034097 | response to cytokine | 0.0001053 |
| GO:0006987 | activation of signaling protein activity involved in unfolded protein response | 0.0001053 |
| GO:0071499 | cellular response to laminar fluid shear stress | 0.0001053 |
| GO:0000398 | mRNA splicing, via spliceosome | 0.0001095 |
| GO:0048705 | skeletal system morphogenesis | 0.0001226 |
| GO:0071353 | cellular response to interleukin-4 | 0.0001237 |
| GO:0016070 | RNA metabolic process | 0.0001291 |
| GO:0043433 | negative regulation of sequence-specific DNA binding transcription factor activity | 0.0001513 |
| GO:0002486 | antigen processing and presentation of endogenous peptide antigen via MHC class I via ER pathway, TAP-independent | 0.000158 |
| GO:0045892 | negative regulation of transcription, DNA-templated | 0.0001617 |
| GO:0048662 | negative regulation of smooth muscle cell proliferation | 0.0002132 |
| GO:0008380 | RNA splicing | 0.0002219 |
| GO:0043434 | response to peptide hormone | 0.0002296 |
| GO:0021762 | substantia nigra development | 0.0002719 |
| GO:0034976 | response to endoplasmic reticulum stress | 0.00029 |

**ADNPs GO Terms**

| **GOID** | **GOTerm** | **P-Value** |
| --- | --- | --- |
| GO:0030198 | extracellular matrix organization | 1.294E-17 |
| GO:0007155 | cell adhesion | 2.035E-10 |
| GO:0030199 | collagen fibril organization | 2.478E-09 |
| GO:0001501 | skeletal system development | 8.734E-09 |
| GO:0045944 | positive regulation of transcription from RNA polymerase II promoter | 1.663E-08 |
| GO:0022617 | extracellular matrix disassembly | 3.188E-07 |
| GO:0000122 | negative regulation of transcription from RNA polymerase II promoter | 6.322E-07 |
| GO:0030574 | collagen catabolic process | 1.183E-06 |
| GO:0007050 | cell cycle arrest | 1.66E-06 |
| GO:0035987 | endodermal cell differentiation | 7.714E-06 |
| GO:0007179 | transforming growth factor beta receptor signaling pathway | 8.171E-06 |
| GO:0001525 | angiogenesis | 1.4E-05 |
| GO:0001568 | blood vessel development | 1.842E-05 |
| GO:0006865 | amino acid transport | 2.386E-05 |
| GO:0030512 | negative regulation of transforming growth factor beta receptor signaling pathway | 3.834E-05 |
| GO:0045444 | fat cell differentiation | 4.298E-05 |
| GO:0048146 | positive regulation of fibroblast proliferation | 5.614E-05 |
| GO:0006366 | transcription from RNA polymerase II promoter | 5.619E-05 |
| GO:0043001 | Golgi to plasma membrane protein transport | 6.09E-05 |
| GO:0003007 | heart morphogenesis | 6.397E-05 |
| GO:0014912 | negative regulation of smooth muscle cell migration | 6.773E-05 |
| GO:0003333 | amino acid transmembrane transport | 7.054E-05 |
| GO:1902966 | positive regulation of protein localization to early endosome | 7.632E-05 |
| GO:0001503 | ossification | 0.000107 |
| GO:0045599 | negative regulation of fat cell differentiation | 0.0001124 |
| GO:0048705 | skeletal system morphogenesis | 0.0001124 |
| GO:0032956 | regulation of actin cytoskeleton organization | 0.00013 |
| GO:0046825 | regulation of protein export from nucleus | 0.0001504 |
| GO:0061028 | establishment of endothelial barrier | 0.0001781 |
| GO:0043277 | apoptotic cell clearance | 0.0001781 |
| GO:0007219 | Notch signaling pathway | 0.0001879 |
| GO:0010628 | positive regulation of gene expression | 0.0002892 |
| GO:0050919 | negative chemotaxis | 0.000301 |
| GO:0048008 | platelet-derived growth factor receptor signaling pathway | 0.0003392 |
| GO:0008284 | positive regulation of cell proliferation | 0.0003679 |
| GO:0048593 | camera-type eye morphogenesis | 0.0003809 |
| GO:0060325 | face morphogenesis | 0.0003956 |
| GO:0035989 | tendon development | 0.0003967 |
| GO:0071709 | membrane assembly | 0.0003967 |
| GO:1903225 | negative regulation of endodermal cell differentiation | 0.0003967 |
| GO:0030335 | positive regulation of cell migration | 0.0003991 |
| GO:0002053 | positive regulation of mesenchymal cell proliferation | 0.0004587 |
| GO:0030154 | cell differentiation | 0.0004762 |
| GO:0060021 | palate development | 0.0004797 |
| GO:0007229 | integrin-mediated signaling pathway | 0.0005359 |
| GO:0045893 | positive regulation of transcription, DNA-templated | 0.0005761 |
| GO:0042340 | keratan sulfate catabolic process | 0.0006041 |
| GO:0051216 | cartilage development | 0.0006198 |

**RegNPs GO Terms**

| **GOID** | **GOTerm** | **P-Value** |
| --- | --- | --- |
| GO:0006954 | inflammatory response | 4.936E-13 |
| GO:0071294 | cellular response to zinc ion | 3.004E-09 |
| GO:0045926 | negative regulation of growth | 9.96E-09 |
| GO:0035666 | TRIF-dependent toll-like receptor signaling pathway | 2E-08 |
| GO:0071222 | cellular response to lipopolysaccharide | 2.027E-08 |
| GO:0002756 | MyD88-independent toll-like receptor signaling pathway | 2.248E-08 |
| GO:0034138 | toll-like receptor 3 signaling pathway | 3.161E-08 |
| GO:0034142 | toll-like receptor 4 signaling pathway | 1.434E-07 |
| GO:0071276 | cellular response to cadmium ion | 1.933E-07 |
| GO:0032496 | response to lipopolysaccharide | 3.658E-07 |
| GO:0002224 | toll-like receptor signaling pathway | 4.694E-07 |
| GO:0010628 | positive regulation of gene expression | 5.726E-07 |
| GO:0034166 | toll-like receptor 10 signaling pathway | 1.709E-06 |
| GO:0034146 | toll-like receptor 5 signaling pathway | 1.709E-06 |
| GO:0048711 | positive regulation of astrocyte differentiation | 1.771E-06 |
| GO:0008285 | negative regulation of cell proliferation | 1.972E-06 |
| GO:0038124 | toll-like receptor TLR6:TLR2 signaling pathway | 3.095E-06 |
| GO:0038123 | toll-like receptor TLR1:TLR2 signaling pathway | 3.095E-06 |
| GO:0034162 | toll-like receptor 9 signaling pathway | 3.398E-06 |
| GO:0034134 | toll-like receptor 2 signaling pathway | 3.726E-06 |
| GO:0006915 | apoptotic process | 3.786E-06 |
| GO:0070374 | positive regulation of ERK1 and ERK2 cascade | 4.314E-06 |
| GO:0070301 | cellular response to hydrogen peroxide | 5.255E-06 |
| GO:0002755 | MyD88-dependent toll-like receptor signaling pathway | 6.844E-06 |
| GO:0071356 | cellular response to tumor necrosis factor | 1.02E-05 |
| GO:0014070 | response to organic cyclic compound | 1.168E-05 |
| GO:0048661 | positive regulation of smooth muscle cell proliferation | 1.242E-05 |
| GO:0071347 | cellular response to interleukin-1 | 1.37E-05 |
| GO:0009612 | response to mechanical stimulus | 1.508E-05 |
| GO:0045600 | positive regulation of fat cell differentiation | 2.168E-05 |
| GO:0043066 | negative regulation of apoptotic process | 2.473E-05 |
| GO:0033138 | positive regulation of peptidyl-serine phosphorylation | 3.34E-05 |
| GO:0007249 | I-kappaB kinase/NF-kappaB signaling | 3.948E-05 |
| GO:0006955 | immune response | 4.057E-05 |
| GO:0045944 | positive regulation of transcription from RNA polymerase II promoter | 4.431E-05 |
| GO:0070373 | negative regulation of ERK1 and ERK2 cascade | 4.914E-05 |
| GO:0043123 | positive regulation of I-kappaB kinase/NF-kappaB signaling | 4.956E-05 |
| GO:0060546 | negative regulation of necroptotic process | 5.314E-05 |
| GO:2000116 | regulation of cysteine-type endopeptidase activity | 7.62E-05 |
| GO:0002246 | wound healing involved in inflammatory response | 7.62E-05 |
| GO:0039535 | regulation of RIG-I signaling pathway | 7.62E-05 |
| GO:0036018 | cellular response to erythropoietin | 7.62E-05 |
| GO:0070427 | nucleotide-binding oligomerization domain containing 1 signaling pathway | 7.62E-05 |
| GO:0002237 | response to molecule of bacterial origin | 0.000103 |
| GO:0051403 | stress-activated MAPK cascade | 0.0001277 |
| GO:0045651 | positive regulation of macrophage differentiation | 0.0001365 |
| GO:0070555 | response to interleukin-1 | 0.0001673 |
| GO:0010243 | response to organonitrogen compound | 0.0001673 |
| GO:0032743 | positive regulation of interleukin-2 production | 0.0001763 |
| GO:0006953 | acute-phase response | 0.0002129 |

**Table 4 Comparsion of HT-CLNPs-I and HT-CLNPs-II**

**HT-CLNPs-I GO Terms**

| **GOID** | **GOTerm** | **P-Value** |
| --- | --- | --- |
| GO:0006048 | UDP-N-acetylglucosamine biosynthetic process | 3.524E-07 |
| GO:0012501 | programmed cell death | 1.952E-06 |
| GO:0008285 | negative regulation of cell proliferation | 2.502E-06 |
| GO:0042026 | protein refolding | 3.032E-06 |
| GO:0010467 | gene expression | 1.394E-05 |
| GO:0043066 | negative regulation of apoptotic process | 1.964E-05 |
| GO:0031397 | negative regulation of protein ubiquitination | 2.025E-05 |
| GO:0010941 | regulation of cell death | 2.151E-05 |
| GO:0007596 | blood coagulation | 2.385E-05 |
| GO:1900740 | positive regulation of protein insertion into mitochondrial membrane involved in apoptotic signaling pathway | 4.689E-05 |
| GO:0045648 | positive regulation of erythrocyte differentiation | 4.689E-05 |
| GO:0002042 | cell migration involved in sprouting angiogenesis | 6.276E-05 |
| GO:1900034 | regulation of cellular response to heat | 7.543E-05 |
| GO:0006047 | UDP-N-acetylglucosamine metabolic process | 8.473E-05 |
| GO:0048010 | vascular endothelial growth factor receptor signaling pathway | 9.284E-05 |
| GO:0045944 | positive regulation of transcription from RNA polymerase II promoter | 0.0001026 |
| GO:0006928 | cellular component movement | 0.0001158 |
| GO:0006915 | apoptotic process | 0.0001216 |
| GO:0034605 | cellular response to heat | 0.0001572 |
| GO:0007623 | circadian rhythm | 0.0001957 |
| GO:0032092 | positive regulation of protein binding | 0.0002012 |
| GO:0042267 | natural killer cell mediated cytotoxicity | 0.0002278 |
| GO:0034629 | cellular protein complex localization | 0.0002315 |
| GO:0033173 | calcineurin-NFAT signaling cascade | 0.0002315 |
| GO:0045893 | positive regulation of transcription, DNA-templated | 0.0002562 |
| GO:0007264 | small GTPase mediated signal transduction | 0.0002668 |
| GO:0006986 | response to unfolded protein | 0.0003037 |
| GO:0032060 | bleb assembly | 0.000343 |
| GO:0006357 | regulation of transcription from RNA polymerase II promoter | 0.0003793 |
| GO:0097193 | intrinsic apoptotic signaling pathway | 0.0004046 |
| GO:0006367 | transcription initiation from RNA polymerase II promoter | 0.0004339 |
| GO:0050821 | protein stabilization | 0.0004345 |
| GO:0010628 | positive regulation of gene expression | 0.0004551 |
| GO:0061045 | negative regulation of wound healing | 0.0004841 |
| GO:0070374 | positive regulation of ERK1 and ERK2 cascade | 0.000569 |
| GO:0051085 | chaperone mediated protein folding requiring cofactor | 0.0006576 |
| GO:0006366 | transcription from RNA polymerase II promoter | 0.0006606 |
| GO:0045597 | positive regulation of cell differentiation | 0.0007432 |
| GO:0060128 | corticotropin hormone secreting cell differentiation | 0.0007994 |
| GO:0070370 | cellular heat acclimation | 0.0007994 |
| GO:2000544 | regulation of endothelial cell chemotaxis to fibroblast growth factor | 0.0007994 |
| GO:0043123 | positive regulation of I-kappaB kinase/NF-kappaB signaling | 0.0008046 |
| GO:0033138 | positive regulation of peptidyl-serine phosphorylation | 0.0010132 |
| GO:0060129 | thyroid-stimulating hormone-secreting cell differentiation | 0.0015814 |
| GO:0060591 | chondroblast differentiation | 0.0015814 |
| GO:0010664 | negative regulation of striated muscle cell apoptotic process | 0.0015814 |
| GO:0022614 | membrane to membrane docking | 0.0015814 |
| GO:0015936 | coenzyme A metabolic process | 0.0015814 |
| GO:0070434 | positive regulation of nucleotide-binding oligomerization domain containing 2 signaling pathway | 0.0015814 |
| GO:0045765 | regulation of angiogenesis | 0.0015841 |

**HT-CLNP-II GO Terms**

| **GOID** | **GOTerm** | **P-Value** |
| --- | --- | --- |
| GO:0030198 | extracellular matrix organization | 8.868E-15 |
| GO:0022617 | extracellular matrix disassembly | 9.881E-15 |
| GO:0006614 | SRP-dependent cotranslational protein targeting to membrane | 1.733E-14 |
| GO:0006414 | translational elongation | 1.073E-13 |
| GO:0030574 | collagen catabolic process | 1.158E-13 |
| GO:0006415 | translational termination | 5.037E-13 |
| GO:0071294 | cellular response to zinc ion | 7.805E-13 |
| GO:0019083 | viral transcription | 7.948E-12 |
| GO:0000184 | nuclear-transcribed mRNA catabolic process, nonsense-mediated decay | 2.552E-11 |
| GO:0019058 | viral life cycle | 2.9E-10 |
| GO:0045926 | negative regulation of growth | 3.204E-10 |
| GO:0006413 | translational initiation | 8.987E-10 |
| GO:0016071 | mRNA metabolic process | 9.684E-10 |
| GO:0030199 | collagen fibril organization | 2.477E-09 |
| GO:0016070 | RNA metabolic process | 4.614E-09 |
| GO:0071276 | cellular response to cadmium ion | 5.567E-09 |
| GO:0001501 | skeletal system development | 2.605E-08 |
| GO:0006412 | translation | 4.843E-08 |
| GO:0016032 | viral process | 2.739E-07 |
| GO:0070208 | protein heterotrimerization | 1.033E-06 |
| GO:0009612 | response to mechanical stimulus | 1.974E-06 |
| GO:0071230 | cellular response to amino acid stimulus | 3.416E-06 |
| GO:0007568 | aging | 3.571E-06 |
| GO:0044267 | cellular protein metabolic process | 3.72E-06 |
| GO:0001957 | intramembranous ossification | 1.753E-05 |
| GO:0010467 | gene expression | 2.154E-05 |
| GO:0030168 | platelet activation | 2.648E-05 |
| GO:0007179 | transforming growth factor beta receptor signaling pathway | 3.104E-05 |
| GO:0009629 | response to gravity | 4.839E-05 |
| GO:0007566 | embryo implantation | 5.748E-05 |
| GO:0036018 | cellular response to erythropoietin | 9.356E-05 |
| GO:1903225 | negative regulation of endodermal cell differentiation | 9.356E-05 |
| GO:0035989 | tendon development | 9.356E-05 |
| GO:0043206 | extracellular fibril organization | 0.0001396 |
| GO:0043065 | positive regulation of apoptotic process | 0.0001475 |
| GO:0060707 | trophoblast giant cell differentiation | 0.0001848 |
| GO:0043392 | negative regulation of DNA binding | 0.0002478 |
| GO:0000302 | response to reactive oxygen species | 0.0002478 |
| GO:0032570 | response to progesterone | 0.0002799 |
| GO:0034097 | response to cytokine | 0.0002857 |
| GO:0001666 | response to hypoxia | 0.0003199 |
| GO:0007623 | circadian rhythm | 0.0003605 |
| GO:0008284 | positive regulation of cell proliferation | 0.0005321 |
| GO:0048705 | skeletal system morphogenesis | 0.0005377 |
| GO:0048592 | eye morphogenesis | 0.0005542 |
| GO:0097084 | vascular smooth muscle cell development | 0.0005542 |
| GO:0032374 | regulation of cholesterol transport | 0.0005542 |
| GO:0008283 | cell proliferation | 0.0006918 |
| GO:0007155 | cell adhesion | 0.0007773 |
| GO:0006898 | receptor-mediated endocytosis | 0.0008835 |
